# How long do Red Queen dynamics survive under genetic drift? A comparative analysis of evolutionary and eco-evolutionary models

**DOI:** 10.1101/490201

**Authors:** Hanna Schenk, Hinrich Schulenburg, Arne Traulsen

## Abstract

**Background:** Red Queen dynamics are defined as long term co-evolutionary dynamics, often with oscillations of genotype abundances driven by fluctuating selection in host-parasite systems. Much of our current understanding of these dynamics is based on theoretical concepts explored in mathematical models that are mostly (i) deterministic, inferring an infinite population size and (ii) evolutionary, thus ecological interactions that change population sizes are excluded. Here, we recall the different mathematical approaches used in the current literature on Red Queen dynamics. We then compare models from game theory (evo) and classical theoretical ecology models (eco-evo), that are all derived from individual interactions and are thus intrinsically stochastic. We assess the influence of this stochasticity through the time to the first loss of a genotype within a host or parasite population.

**Results:** The time until the first genotype is lost (“extinction time”), is shorter when ecological dynamics, in the form of a changing population size, is considered. Furthermore, when individuals compete only locally with other individuals extinction is even faster. On the other hand, evolutionary models with a fixed population size and competition on the scale of the whole population prolong extinction and therefore stabilise the oscillations. The stabilising properties of intraspecific competitions become stronger when population size is increased and the deterministic part of the dynamics gain influence. In general, the loss of genotype diversity can be counteracted with mutations (or recombination), which then allow the populations to recurrently undergo negative frequency-dependent selection dynamics and selective sweeps.

**Conclusion:** Although the models we investigated are equal in their biological motivation and interpretation, they have diverging mathematical properties both in the derived deterministic dynamics and the derived stochastic dynamics. We find that models that do not consider intraspecific competition and that include ecological dynamics by letting the population size vary, lose genotypes – and thus Red Queen oscillations – faster than models with competition and a fixed population size.

## Background

Diversity, induced by continuous co-evolution can theoretically be maintained by the intense antagonistic relationship of hosts and parasites. This is the central part of the Red Queen hypothesis, verbally first formulated by van Valen in 1973 [1]. The hypothesis has been mathematically formulated in many models. However, owing to the modern usage of the term ‘Red Queen’ for different but related phenomena [2, 3, 4, 5, 6, 7, 8, 9], the models have diverging foci and many lack the implementation of stochastic forces and ecological dynamics. A common synonym for the term Red Queen dynamics is fluctuating selection dynamics (FSD). Such fluctuations can be induced by co-evolving hosts and parasites and, as one possibility, be driven by negative frequency-dependent selection (NFDS), where host and parasite genotype abundances oscillate in time and every genotype can temporally be best adapted. In detail, since parasites are selected to target the most common resource, being a rare host genotype is advantageous. This temporary high fitness makes the genotype grow in relative abundance, but before it can take over the whole population, it is severely diminished by the profiting parasites genotypes, which target this now common host type. By contrast, in arms race dynamics (ARD) novel favoured genotypes spread in the entire population by recurrent selective sweeps. The terms NFDS and ARD are both referred to as Red Queen dynamics [10, 11, 12] and describe an ongoing co-evolutionary change without approaching an equilibrium. In this paper, we use the term Red Queen dynamics for NFDS, as is commonly done in the literature, but return to other definitions of the Red Queen in the discussion.

Although Red Queen dynamics is a well-known and frequently cited concept, there is only little evidence for the ubiquitous prevalence of long term Red Queen dynamics in nature – empirical challenges preclude the observation of more than a few subsequent oscillations, as these require a major amounts of intensive and challenging lab work [13, 14, 15, 16, 17]. Thus, most work on the actual long term temporal dynamics is theoretical, often dealing with evolutionary dynamics or epidemiological dynamics in a deterministic fashion. We have summarised some of the literature in the context of these assumptions in Table 1 (methods in the additional file). Similar literature summaries exist with a focus on sexual vs. asexual reproduction [8] or host-parasite coevolution models [64]. Many theoretical studies build on evolutionary game theory [19] and a zero-sum assumption, where the harm done to the host equals the benefit for the parasite, which was already envisioned by van Valen at the time. Some of the models are implemented with equations that describe both species’ dynamics (explicit host-parasite HP dynamics), other studies, especially on the evolution and maintenance of sexual reproduction (Red Queen Hypothesis) revert to epidemiological models (susceptible-infected SI models), sometimes in the pursuit of including population dynamics. The present work focuses on evolutionary host-parasite models in comparison with eco-evolutionary models that include population dynamics without using SI models.

**Table 1: -.**
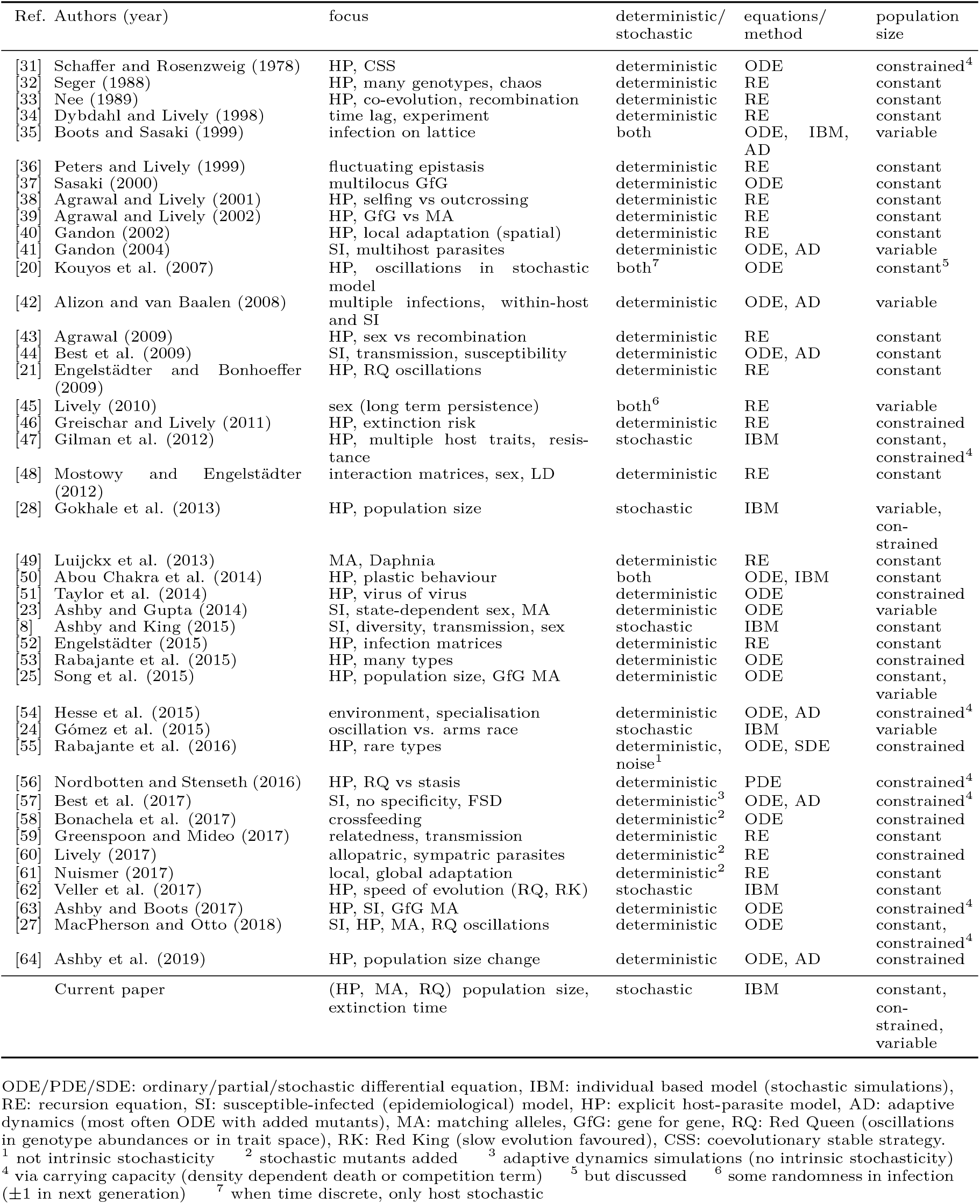
Literature overview. Mathematical models and properties discussed in this paper sorted by publication year. Many models deal with relative allele or genotype abundances without considering ecological dynamics – these have been categorised as constant population size models. Those models that do include a changing population size and stochastic effects mostly do not analyse the stability of long term oscillations which is the focus of this paper. (See the notes on this literature survey in the additional file).

While many studies assess the occurrence of oscillating selection dynamics and show under what assumptions oscillations dominate [64, 20, 21, 22, 23, 24, 25, 26, 27], only few studies include both ecological population dynamics and stochastic noise, although the combination of the two has been shown to result in a fast loss of genotypes in either population [28]. It has been difficult to derive a stochastic model that easily switches between constant and changing population size using a single parameter. For example Gokhale et al. [28] artificially normalised population size every few generations. Here, we take a different approach and compare the modelling framework of evolutionary game theory, where population size is constant by design, to eco-evolutionary dynamics from the field of theoretical ecology, where population size is inherently free to change over time. Our goal is not to present the one model that is the best description of reality, but to illustrate how different modelling assumptions can drive the results from such models.

Specifically, we use individual-based models, since ecological and evolutionary dynamics of populations are driven by events on the individual level. The models are based on haploid and asexual populations that live in a well-mixed environment where encounters are density dependent. Individuals are born, interact with other individuals of their own or opposing species and die. Generally, we will consider at least two genotypes and track the associated abundances *H*_1_, *H*_2_, *P*_1_, *P*_2_ and the total population sizes *N_H_*, *N_P_* of hosts and parasites over time. We simulate the dynamics using a uniformly distributed initial standing genetic variation and the simple matching allele interaction profile, where parasites are highly specialised [29, 30] on a particular host genotype and identical in all other aspects. Yet, the way this interaction profile enters in the dynamical equations and thus defines fitness for the individual genotypes is very different between the models. In population dynamics models these events happen at constant rates and depending on the density of the interacting individuals. A similarly simple, yet completely different approach is the stochastic birth-death process which tracks only the evolutionary dynamics. In each time step one individual is born, proportional to its current ‘fitness’ and another individual dies proportional to its density.

These models all produce Red Queen dynamics (NFDS) and we assess the robustness of those by measuring the time to extinction, which we define as the earliest time that any genotype from the initial genetic variation is lost in either population. Further, we consider the impact of the derived deterministic dynamics and the influence of ecology in the form of a population-size-change on this extinction time. The time to extinction of a genotype represents the durability of the stochastic oscillations. Without the immigration or re-emergence of extinct genotypes, the diversity of both populations declines in the long run.

## Results

Evolutionary dynamics depict the change of relative genotype abundances over time and can be examined without keeping track of population size changes. However, it is well known that ecological dynamics can feed back on evolutionary dynamics. We want to understand this feedback in the context of Red Queen dynamics. To this end we compare models from evolutionary game theory, that do not include population size changes and theoretical ecology models that do. The models have been widely used in the literature and represent the simplest case of Red Queen dynamics with a matching allele interaction profile (for details see the Methods below and additional file).

### 1. The matching-allele host-parasite Red Queen dynamics in evolutionary and eco-evolutionary models

In an evolutionary birth-death process one individual is born and another dies in each population, here host or parasite, and in each time step. Thereby, population size remains constant and the focus lies on the genotypic composition of a population. The Evo^+^ and Evo processes (see Table 2 and Methods for a definition) are such birth-death processes [65, 66, and references therein]. Individuals are chosen to die randomly, but the individual that reproduces is chosen proportional to the fitness advantages of that genotype relative to other genotypes in the population. The fitness effects are imposed by the current state of the antagonist population and an interaction matrix. In the Evo^+^ process, the fitness effect is normalised by the average fitness effect over the whole host population, which leads to a kind of intra-specific competition (+) while in the Evo process the difference in fitness effects is compared between a genotype-one individual and a genotype-two individual, thus competition is pairwise. Because of the population size constraint, both processes can be analytically treated (see additional file) when implemented in discrete time (prefix dt).

**Table 2: -.** Model overview. Model names and their main differences. The Evo^+^ and Evo model are derived from evolutionary game theory while the EcoEvo^+^ and EcoEvo model stem from theoretical ecology. The Hybrid model combines elements from both. Models are ordered by population size constraint. The deterministic dynamics apply to the two types matching alleles interaction matrix. Details on the models and analysis are available in the additional file.

| | Model | description | features | stochastic dynamics (small $N$)<sup>×</sup> | deterministic dynamics (large $N$)<sup>*</sup> | stochastic dynamics (medium $N$)<sup>†</sup> | population size change |
| --- | --- | --- | --- | --- | --- | --- | --- |
| evolution. game theory | $\text{Evo}^+$ | Birth-death process. Which individual reproduces depends on the current fitness effect by the antagonist, normalised by the population average fitness effects (+) | intraspecific competition (+) | slow extinction | stasis | NFDS | no |
| | $\text{Evo}$ | Like $\text{Evo}^+$ but fitness effects are compared between two individuals not with the population average | pairwise competition | slow extinction | NFDS | extinction | no |
|  | Hybrid | Hybrid model with reactions between two genotypes of different populations, single birth of parasite and death of host by dynamically adjusted rates. | no competition | extinction | NFDS | extinction | yes, but dynamically constrained |
| theoret. ecology | $\text{EcoEvo}^+$ | Independent reactions between individual hosts and parasites, single birth and death events or competition in hosts | intra-host competition (+) | fast extinction | stasis | NFDS | yes, but carrying capacity |
| | $\text{EcoEvo}$ | Like $\text{EcoEvo}^+$ but without competition within hosts. For infinite population size this is the Lotka-Volterra dynamics | no competition | fast extinction | NFDS | extinction | yes, unconstrained |

<sup>×</sup> population size *change* speeds up the extinction of genotypes (Fig. 1)
<sup>\*</sup> for large population sizes $N$ the deterministic dynamics dominate (Fig. 2). Damped oscillations lead to an attractive equilibrium ('stasis'). When the equilibrium is neutral genotype abundances oscillate induced by negative frequency-dependent selection ('NFDS').
<sup>†</sup> when population size is intermediate dynamics are strongly influenced by their deterministic characteristics but with stochastic noise. Stochastic dynamics with oscillations are stabilised by the attractive deterministic fixed point which can countervail the stochastic outward pull, postponing extinction ('NFDS'). Without the attractive pull the time to the first extinction of a genotype is much shorter ('extinction').

In models adapted from theoretical ecology the events of birth, death and interaction happen independently with external rates and, importantly, between populations (EcoEvo, comparable with the Lotka-Volterra dynamics in [28]). Host and parasite individuals encounter one another based on their densities and if they match, an interaction is carried out with a constant rate upon which a host dies or a parasite reproduces. When competition between hosts (+) is included, the host population grows logistically with a carrying capacity *K*. The host population size *N_H_* reaches the carrying capacity in the absence of parasites. However, in the presence of parasites the population size is smaller than *K* because of the additional mortality from parasites (see Table 3 for details on the parameters in all models).

**Table 3: -.** Model parameters.

| Parameter | interpretation | default value / range | Models |
| --- | --- | --- | --- |
| $H_i$ | absolute abundances of host genotype $i$ | $\in [0, N_H]$ | all |
| $P_i$ | absolute abundances of parasite genotype $i$ | $\in [0, N_P]$ | all |
| $h_i$ | relative abundances of host genotype $i$ | $\in [0, 1]$ | all |
| $p_i$ | relative abundances of parasite genotype $i$ | $\in [0, 1]$ | all |
| $N_H$ | total host population size $\sum_i H_i$ | - | all |
| $N_P$ | total parasite population size $\sum_i P_i$ | - | all |
| $b_H$ | host birth rate | - | EcoEvo<sup>+</sup>, EcoEvo |
| $d_P$ | parasite death rate | 1 | EcoEvo<sup>+</sup>, EcoEvo |
| $\lambda$ | rate of host death or parasite birth when genotypes match | - | EcoEvo<sup>+</sup>, EcoEvo |
| $\mu$ | intraspecific competition rate | - | EcoEvo<sup>+</sup> |
| $K$ | carrying capacity (host population size in the absence of parasites) | - | EcoEvo<sup>+</sup> |
| $\alpha$ | fitness gain (loss) of matching genotypes for parasite (host) | 1 | Evo<sup>+</sup>, Evo, Hybrid |
| $\beta$ | fitness gain (loss) of mismatching genotypes for parasite (host) | 0 | Evo<sup>+</sup>, Evo, Hybrid |
| $w_H, w_P$ | selection intensity | $\in [0, 1]$ | Evo<sup>+</sup>, Evo, Hybrid |
| $\pi_{H_i}, \pi_{P_i}$ | payoff from the game, defined for each genotype | - | Evo<sup>+</sup>, Evo, Hybrid |
| $\underline{f}_{H_i}, \underline{f}_{P_i}$ | ‘fitness’ from the game, defined for each genotype | - | Evo<sup>+</sup>, Evo, Hybrid |
| $\bar{f}_H, \bar{f}_P$ | average ‘fitness’, defined for each population | - | Evo<sup>+</sup>, Evo, Hybrid |
| $d_{H_i}$ | death rate of host genotype $i$ | - | Hybrid |
| $b_{P_i}$ | birth rate of parasite genotype $i$ | - | Hybrid |

Both Evo and EcoEvo modelling approaches are combined in an intermediate model with self-controlled, but not fixed, population size (Hybrid). The model is implemented as an individual interactions model, where reactions take place also between populations, but the rates of these events are taken from the game theory models: Host death and parasite birth happen according to the fitness effects, host death and parasite birth rates are then adapted dynamically to keep population size nearly constant.

From the derivations of the models (details in Methods and additional file) some basic properties of the dynamics are obtained and summarised in Table 2. The evolutionary game theory models have a constant population size by design, whereas population size can change in all other models. The average behaviour of the individual-based stochastic processes is captured in the deterministic selection term and the noise term, which together determine the stochastic dynamics. The noise term is discussed in Section 2 (Figure 1). The role of intra-specific competition in the deterministic part is discussed in Section 3 (Figure 2).

**Figure 1: -.**
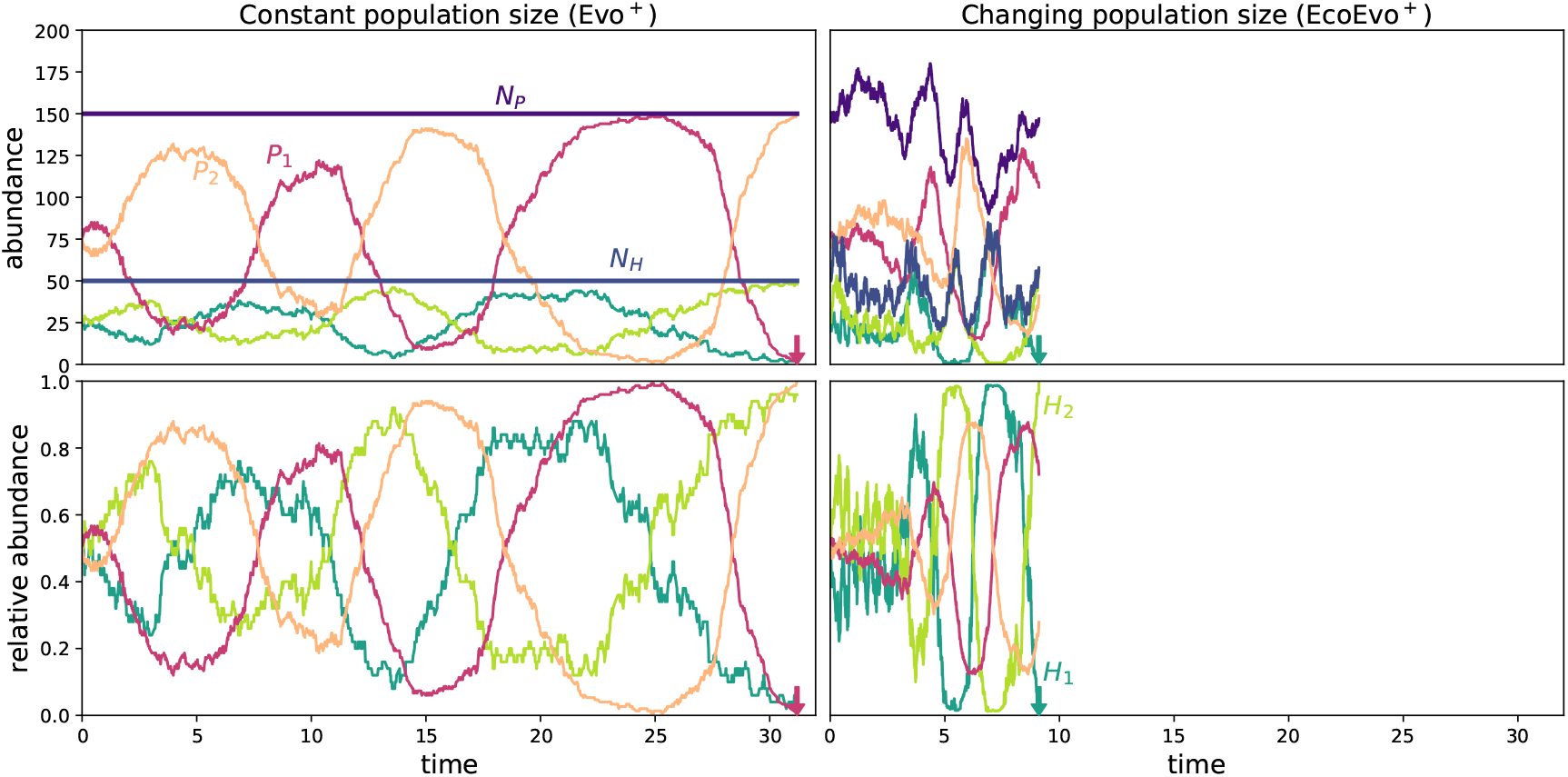
Example run illustrating that extinction is faster with ecological dynamics. Oscillations of host and parasite genotype abundances in the Evo^+^ process with constant population size and EcoEvo^+^ process with changing population size. The simulations start with an equal abundance of both genotypes *H*_1_(0) = *H*_2_(0) = *N_H_*/2 and *P*_1_(0) = *P*_2_(0) = *N_P_*/2. Method: Simulation of the stochastic processes with the Gillespie algorithm. Parameters: Total population sizes *N_H_* = 50, *N_P_* = 150 (only initially for the EcoEvo^+^ model), selection strengths *w_H_* = 0.5, *w_P_* = 1, matching allele parameters *α* = 1, *β* = 0, death rate of the parasite *d_P_* = 1, birth rate of the host *b_H_* = 6, carrying capacity *K* = 100, interaction rate λ_0_ = 4, 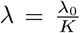, intra-specific competition rate 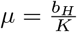. See Methods and additional file for method and parameter details.

**Figure 2: -.**
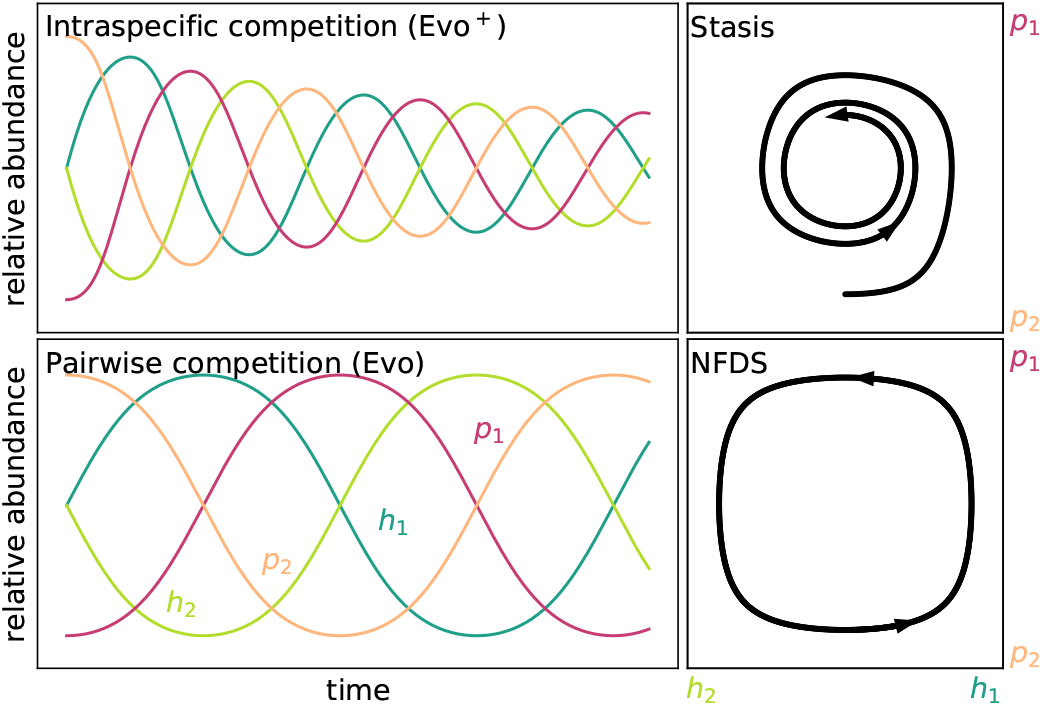
Large population size limit. Relative abundances of two genotypes of host *h*_1_ and *h*_2_ and parasite *p*_1_ and *p*_2_ over time (left) and 2D representation (right) in the deterministic equivalents of the Evo^+^ and Evo process with constant population size. Top: Intraspecific competition within the whole population (+) results in an attracting fixed point which is reached eventually and does not changed once reached, leading to stasis (also EcoEvo^+^). Bottom: Pairwise competition between individuals allows for a neutrally stable fixed point which neither attracts nor repulses the dynamics resulting in continuous co-evolution in the form of negative frequency-dependent selection dynamics (NFDS) around the internal fixed point (also EcoEvo). Method: integration of ordinary differential equations, the adjusted replicator dynamics (Evo^+^) and the replicator dynamics (Evo), which are the deterministic limits of the respective stochastic processes. Parameters: selection strength *w_H_* = *w_P_* = 1, matching allele parameters *α* = 1 and *β* = 0.

### 2. In models with ecological dynamics genotypes die out faster

It is clear that populations with low total population sizes are more prone to genetic drift and the loss of genotypes than large populations. We now show that it is not only the population size but the possibility of population size *change* that speeds up this process. We exemplify our argument here, but provide a more detailed analysis below. As an example with two host genotypes and two parasite genotypes we select the Evo^+^ process and the EcoEvo^+^ process (Figure 1). Starting with an equal abundance of genotypes we measure the time to the first loss of a genotype. When a genotype has died the population becomes monoclonal and oscillations are no longer possible. With the fixed population size in the Evo^+^ process oscillations survive longer than in the EcoEvo^+^ process with a changing population size. The evolutionary dynamics are similar and defined through the relative abundance of the types, but the population size change can speed up the frequency of event occurrences and increase the probability of extinction through the bottleneck effect when population sizes are low.

### 3. Intraspecific competition stabilises negative frequency-dependent selection

The equations that define the stochastic process consist of a deterministic selection term and a noise term and represent the mean and variance of many individual simulations. Therefore, it is impossible to understand the stochastic model without making the deterministic dynamics clear. Furthermore, when population size is large, the stochastic process approaches the more manageable deterministic dynamics (details in the additional file). The deterministic equations for all models from Table 2 have an internal co-existence fixed point, where both genotypes exist in a fixed ratio, which does not change over time. This point is only attractive, if starting with suitable initial compositions of genotypes the dynamics approach the state, in this case in the form of damped oscillations. The intraspecific competition (+) in the Evo^+^ process and the EcoEvo^+^ process result in such an attractive pull (Figure 2). A second possibility is neutral stability, where genotype abundances oscillate with a constant amplitude and period, which depend on the initial abundances. These neutral cycles are produced by models where individuals only compete with other genotypes locally like in the Evo process with pairwise competition or the EcoEvo model with no intraspecific competition and the Hybrid model (Figure 2).

In our models, the noise in the stochastic dynamics always leads to extinction, while deterministic dynamics never do. When population size is large enough to be impacted by the deterministic behaviour but stochastic noise still plays a role, the global competition models (+) show persisting Red Queen oscillations. The deterministic ‘pull’ and the stochastic ‘push’ balance [67], prolonging extinction times. For models with neutral oscillations (NFDS) in the deterministic dynamics stochastic effects will on average increase the amplitudes and push the trajectories to the edges of the space towards a faster extinction of genotypes.

The single simulations (Figure 1) are only a snapshot and one specific realisation of the stochastic processes. Ideally, we would analytically derive general extinction times depending on the parameters of the model. Yet, to derive an exact analytical solution for this problem is extremely challenging. In addition to simulations, we have calculated the numerical (but exact) extinction times for low population sizes and provide an approximative method based on the averaged noise (see additional file for further details). These methods are limited to a subset of the models and can thus not be used for a comparison of all models, but only to support the computationally costly simulations which provide the now following main result.

### 4. The strength of random effects depends on the model properties

We simulate 1000 replicates for a set of parameter combinations of the models with two genotypes in each population and record the time it takes until one genotype has died out. As a general trend, the more constrained a population size is, the longer oscillations survive (higher extinction times in Figure 3). This holds true for small to intermediate population sizes – note a similar vertical order of extinction times to the ordering of models by population size constraint in Table 2. When population sizes become larger and the deterministic model properties gain influence, models with competition terms (+, stasis, compare with Figure 2) have higher extinction times and therefore more stable Red Queen oscillations.

**Figure 3: -.**
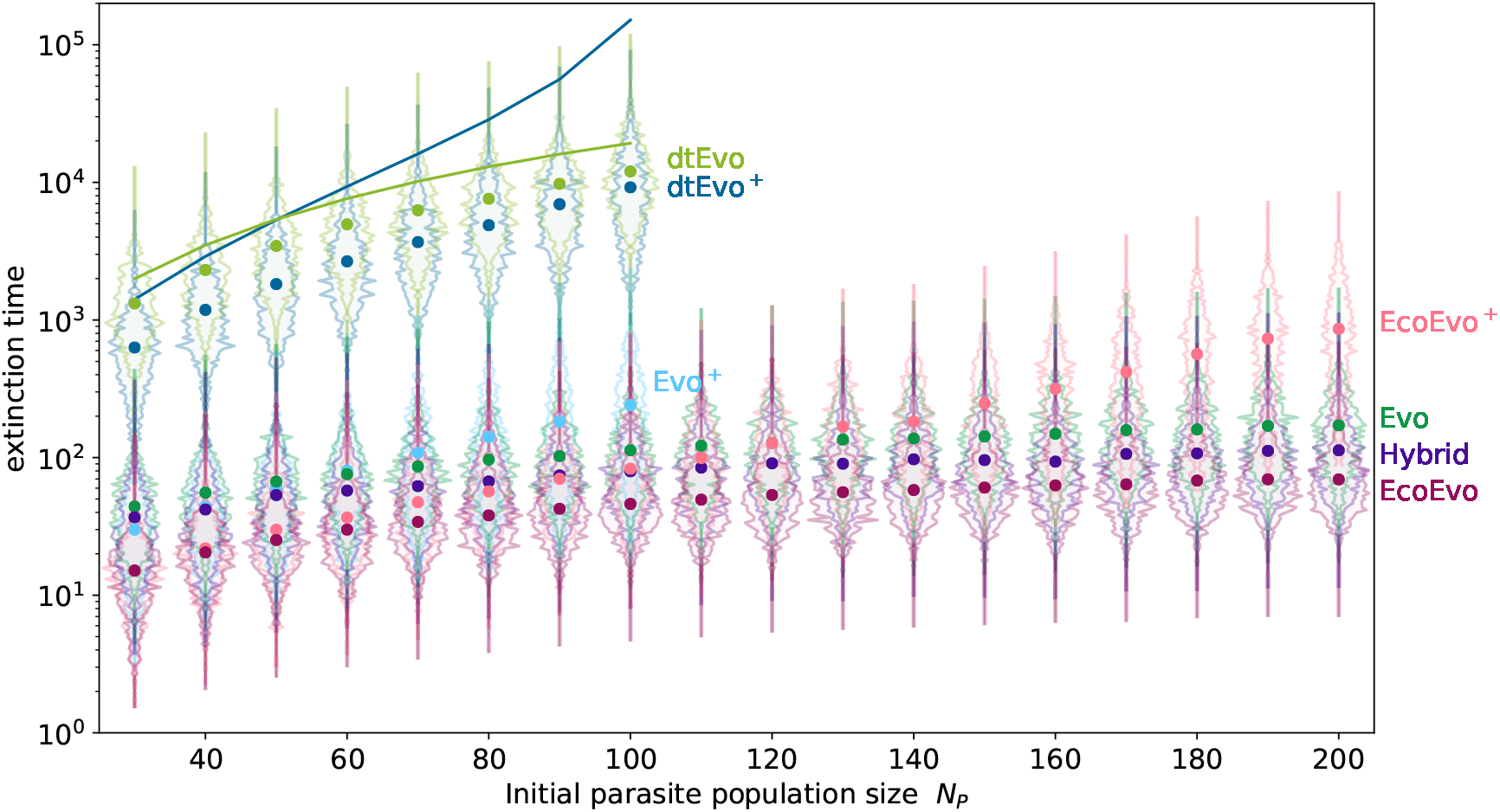
**Extinction time** of either genotype of either host or parasite population for different initial population sizes of the parasite *N_P_* for all models. We show the mean extinction time of any genotype over 1000 independent simulations (fat dots) and the distribution of those extinction times (shaded histogram area around the mean). The simulations start with equal abundance of both genotypes *H*_1_(0) = *H*_2_(0) = *N_H_*/2 and *P*_1_(0) = *P*_2_(0) = *N_P_*/2. Lines denote approximate results based on the average noise (see additional file). The discrete time processes are simulated for values of *N_P_* for which analytical results are valid. The Evo^+^ process is not simulated for high parasite population sizes since the computation becomes extremely time-consuming and the trend is already clear. Parameters as in Figure 1 except *N_H_* = 250, *K* = 500, birthrate *b_h_* ∈ {0.24. 0.32, …, 1.6} in the EcoEvo^+^ and for the EcoEvo model *b_h_* ∈ {0.12. 0.16, …, 0.8} and *μ* = 0 to achieve the population sizes *N_P_* displayed.

By design, the discrete time (dt) processes have much higher extinction times and are thus not directly comparable to the continuous time simulations. A scaling would be possible for equal population sizes, but with different extinction routes and *N_H_* ≠ *N_P_* no such factor can be derived. The dtEvo^+^ and dtEvo extinction times in Figure 3 can therefore only be compared between them. For growing *N_P_*, the dtEvo^+^ process has an increased extinction time because of the stabilising attractive fixed point. This trend is even more pronounced in the approximate analytic solution (solid lines), inspired by Claussen [68, 69] (see additional file). The error of the analytical approach cannot be neglected, but the qualitative trend is clearly visible and the result is fully analytical.

Due to the challenges of employing an exact analytical approach, we cannot perfectly tune the models for the same amplitudes, fluctuations and frequencies/periods of oscillations. The specific choice of the parameters is not necessarily directly comparable, but we have made an effort to choose them in a meaningful way, such that the fixed points are exactly the same and amplitudes comparable. We choose strong selection for the parasite *w_P_* = 1 and weaker selection for the host *w_H_* = 0.5 in the models derived from game theory, because the EcoEvo^+^ model is built in a similar way: Parasite birth can only occur through the antagonistic interaction, but host mortality is also influenced by the competition term. While the parasite is obligate and thus completely dependent on the host, the host suffers, but does not always die from an infection.

The impact of selection intensities on the extinction times is further explored in the additional file. We find that strongly diverging host and parasite selection intensities can counter-intuitively lead to more stable dynamics in the Evo process than in the Evo^+^ process.

### 5. Diversity inflow results in sequential negative frequency-dependent selection dynamics and arms race dynamics

So far we have compared models with two genotypes in each species. We now provide an outlook of how diversity changes for many genotypes. We simulate one possible example with an initial uniform distribution of twenty genotypes in each species (see additional file). Diversity, simply defined as the number of genotypes present in the population, declines exponentially with time at a constant rate. The manual re-introduction of an extinct, but temporarily best adapted parasite genotype can result in a selective sweep that leaves the parasite population monoclonal.

In reality, our genotypes are not as static in their traits as described here, but one of our ‘genotypes’ can actually be seen as an average of several individuals with slightly different traits. We now add a form of mutation or recombination to the model so that reproduction does not necessarily result in a clonal daughter, but a new individual with different traits. For example, parasites could evolve quickly by allowing beneficial mutations to produce other, even extinct, genotypes. Depending on the model system, a sexually reproducing host could also store genetic material to revive long extinct phenotypes by recombination. We abstract this by inserting a conversion rate *μ* from one genotype to the neighbouring genotype. For example with five predefined genotypes we have 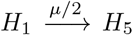 and 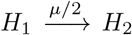 and so on. The dynamics we observe now (Figure 4) are not pure negative frequency-dependent selection dynamics, but a mixture of oscillations and arms race dynamics, where selective sweeps can make a population monoclonal in a very short time, but a re-introduction of extinct genotypes allows for oscillations to re-emerge.

**Figure 4: -.**
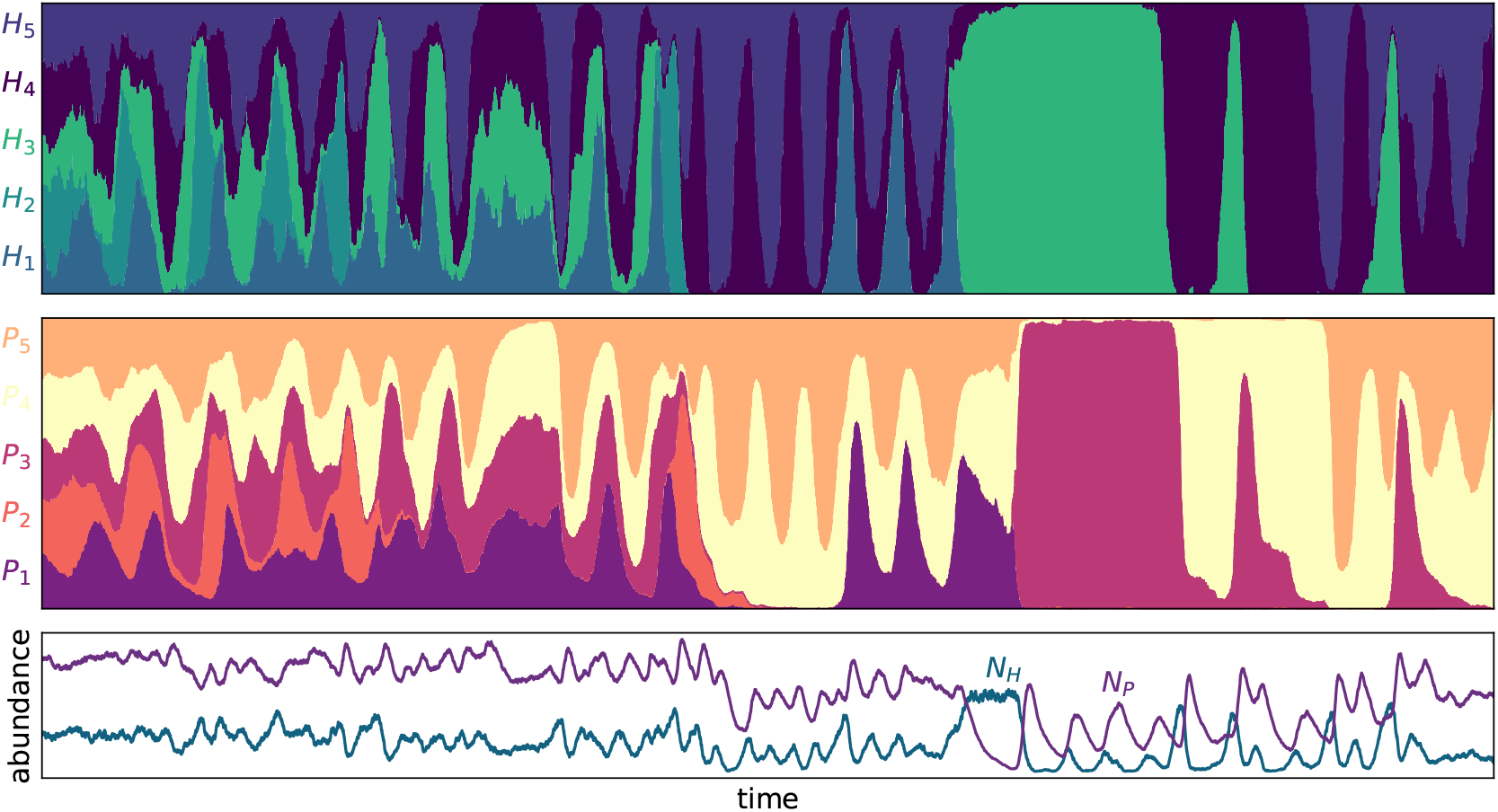
Negative frequency-dependent selection and arms race dynamics. Revival of genotypes and evolution of host (top) and parasite (middle) populations with five possible genotypes each. With the rate *μ_H_* = 0.005 and *μ_P_* = 0.01 genotypes convert to neighbouring genotypes through mutation or recombination. Stacked plots – evolutionary dynamics: the area covered by one colour is proportional to the relative abundance of that genotype of host or parasite at that time. Lower panel – ecological dynamics: total abundance of hosts and parasites. Method: the example is a stochastic simulation (Gillespie algorithm) of an EcoEvo^+^ process. The simulations start with equal abundance of all five genotypes *H_i_*(0) = *N_H_*/5 and *P_i_*(0) = *N_P_*/5 for *i* = 1, 2, 3, 4, 5. Parameters: *N_H_* = 300, *N_P_* = 900 (both initially), *b_H_* =6, *d_P_* = 1, *K* = 600, λ_0_ = 10.

## Discussion

In this paper we compare evolutionary models from evolutionary game theory to eco-evolutionary models from theoretical ecology to understand the impact of ecology and other model properties on the long term co-evolutionary Red Queen oscillations of host and parasite genotypes. The models are individual-based and intrinsically stochastic, thereby allowing genetic drift and the loss of genotypes from a population. Starting with an initially uniform distribution of genotypes, we define the extinction time as the first time that any genotype is lost from any of the two populations, and use this extinction time to measure the robustness of the Red Queen cycles and therefore, the maintenance of diversity. Our main result is that including ecology in models, in the form of a changing population size, leads to a faster loss of genotypes, when stochastic dynamics are considered. This result is similar to the simulation results by Gokhale et al. [28], where ecological dynamics were artificially removed from the simulations, in an attempt to make a straightforward comparison of eco-evo and evo dynamics. In contrast, we compare two modelling frameworks with historically developed differences between them. The models presented here are all based on the same widely used biological assumptions – haploid well-mixed host and parasite genotypes that interact through the matching-alleles infection matrix – but with differences in their mathematical properties: discrete and continuous time models with attractive or neutral deterministic dynamics. The models are further intrinsically stochastic, since they are derived from interactions between individuals. This inherent noise, genetic drift, also impacts the models within a given framework. The mean outward pull by noise that increases amplitudes and thus makes extinction more probable can be counteracted by intraspecific competition that pulls the dynamics back, decreases amplitudes and thus stabilises negative frequency-dependent selection dynamics, resulting in longer extinction times. Finally, we provide a snapshot of what happens when standing genetic variation is large initially. If no inflow of genotypes via mutation or migration is provided, the number of genotypes in a population will decline exponentially. However, when conversions between neighbouring types are allowed with a small mutation rate, negative frequency-dependent selection dynamics and selective sweeps can occur sequentially.

Previous theoretical studies have similarly examined the persistence of Red Queen oscillations under ecological feedbacks. For example Goméz et al. [24] found fluctuating selection and arms race dynamics in an epidemic model (host-focussed) with explicitly modelled parasite populations. MacPherson and Otto [27] also combined epidemiological and neutrally stable host-parasite dynamics and showed that this can dampen allele frequency oscillations, which leads to stasis in their deterministic model but would return to oscillations under stochasticity (see Table 2). Recently, the game theoretical replicator dynamics were mathematically tuned for population size influence using a single parameter [64] resulting in damped oscillations and thus stable polymorphism for both matching-alleles and gene-for-gene infection matrices. While we argue that eco-evolutionary feedbacks increase oscillation amplitudes, Ashby et al. argue that oscillation amplitudes are decreased over time. However, the population dynamics in their model were dampened by a maximal value which resembles our intra-specific competition leading to stasis (Figure 2). Furthermore, their models are deterministic which closer resembles our models when population size is large, where stabilising effects have a larger influence. In the more theoretical literature, it is now well established that assumptions such as population size fluctuations and stochasticity can result in more rapid extinction [70, 71, and many more].

The stabilising property of intra-specific competition is documented in the literature [72]. Intraspecific competition (+) enters in our evolutionary models as part of a genotype’s fitness effect that is compared to the focal population’s average fitness, whereas in the eco-evolutionary models it is implemented as an ecological intra-specific competition term. Both the evolutionary and the ecological implementation of this intra-specific competition stabilise the dynamics and lead to stasis following damped oscillations. The more commonly used host-parasite co-evolution models result in neutrally stable oscillations whereas damped oscillations are often seen as a termination of Red Queen dynamics. Yet, exactly this stasis shows similar oscillation patterns when stochasticity perturbs dynamics away from the stable fixed point (noise induced oscillations [67]). In a stochastic world, pure host-parasite dynamics therefore result in fast extinction, which would only be stabilised by intraspecific competition. For larger population sizes, when the stability of the fixed point gains in importance, the dynamics are pulled more towards the inner equilibrium state, making stochasticity less influential. Thus, only for organisms with large population sizes and good mixing, intraspecific competition would not be necessary for sustained Red Queen oscillations.

Although this study does not explicitly analyse modes of reproduction, our final result shows how reviving extinct genotypes can restore Red Queen dynamics. If parasites can evolve more quickly due to shorter generation times and larger numbers, then hosts are given an advantage by being able to “store” genotypes through recombination. Also, if clonal reproduction accumulates mutations (Muller’s ratchet), this could impact population sizes and sexual reproduction would be even more important [73, 74]. Ashby and King [8] devised a stochastic individual based susceptible-infected model with diploid sexual hosts and showed that high diversity cannot maintain sexual reproduction when parasite transmission rates are low. Although our models are more abstract concerning reproduction, we do explicitly model parasites. If parasite populations are well mixed and diverse with high mutation rates, this can again select for higher diversity through sex, like in [24], where fluctuating selection dynamics, and thus high diversity, is more likely when hosts encounter a diverse parasite population and the disease load is high. Furthermore, our models can include global competition in both species or resource competition in the host, which stabilise the oscillating dynamics. More support for recombination during parasite infection was shown in [48], where hosts could optionally switch between two modes of reproduction. See also [4] for a comprehensive connection to the Red Queen Hypothesis for sexual reproduction.

We have shown that in the same setting and with the exact same parameters sequential occurrences of oscillating selection and arms race dynamics are possible. We show only a snapshot and we do not quantify dynamics as is done in [24], but we find it to be an interesting aspect that the dynamics can occur temporarily in the same simulation, with the same settings and assumptions. The more complete picture could include all possibilities discussed in the Red Queen literature: there can be constant extinction, as suggested by van Valen on a taxonomic level and there can be oscillations and arms race dynamics as suggested by host-parasite interactions and the resulting co-evolution. With our preliminary results we might be going too far if we also justify sexual reproduction, yet, without recombination or mutation, diversity decline is inevitable theoretically.

Our models explore stochasticity under different restrictions of population size, while other modelling aspects are kept relatively plain. In the present work, the infection pattern is restricted to the matching alleles model, and the zero-sum assumption, yet this is necessary for oscillations [21]. Other infectivity patterns that result in Red Queen dynamics have not been examined here. The gene-for-gene infection matrix could show similar results since the oscillations are also neutrally stable, yet including ecological dynamics changes the complexity of the cycles [25]. In general the robustness of those cycles under stochasticity would depend on the details of how different infection mechanisms are translated into mathematical equations. Further limitations are the haploidy of both hosts and parasites and thereby asexual reproduction, the lack of life history or infection history and there is no spatial structure and evolution in the values of resistance or infectiousness. We do, however, briefly explore the effects of including more genotypes and mutation as a means to revive genotypes. There is an increasing effort to openly discuss how verbal models and biological assumptions enter into models [27, 75]. Making the assumptions clear and readily available should be the standard for future publications. For stochastic processes the analogous deterministic dynamics should be stated to provide the reader with a more complete picture of stochastic dynamics.

The model predictions presented here although quite abstract may nevertheless apply to the real world. Bottlenecks are likely more common in natural host-parasite associations [76] than usually assumed and, therefore, the interaction dynamics are likely shaped by genetic drift and, thus, stochastic effects. Eco-evolutionary feedbacks have been confirmed to impact the form of co-evolution in bacteria-virus experiments [77]. Increasing diversity in the parasite or higher exposure lead to a shift from negative frequency-dependent selection to arms race dynamics in two bacteria-phage systems [24, 78]. Oscillations alongside incomplete selective sweeps were recently even documented in a nematode-bacteria interaction [17]. It would now be of particular interest to assess the occurrence of bottlenecks, drift and competition in natural host-parasite associations and relate them to the resulting allele frequency dynamics. Such empirical data would help us to obtain a more general understanding of host-parasite co-evolution and potentially question the importance of sustained Red Queen oscillations in this context.

## Conclusions

We have shown that models equal in their verbal biological description can be quite different in their mathematical details, with great consequences for both deterministic and stochastic dynamics. The loss of genotypes is inevitable in stochastic models without mutation or immigration. This extinction is faster when ecological dynamics are considered in an evolutionary model. Competition between genotypes within a species stabilises the dynamics and slows down extinction thus sustaining Red Queen dynamics. The applicability of models to the real world thus depends on the system of interest and mathematical details should be carefully considered for each particular case study. When bottlenecks, drift and competition are observed, the model needs to be adapted accordingly.

## Methods

The following method descriptions are short explanations of the stochastic processes used in this manuscript. The precise equations and methods of analysis can be found in the additional file. The simulation code is provided at https://github.com/HannaSchenk/ShortLifeRQ.

**The discrete time Evo^+^ process (dtEvo^+^)**, also discrete time Moran process, is a stochastic birth-death process, with a constant population size [79], often used in evolutionary game theory (see for example [80, 81] or [82]). Each birth-death reaction has a reaction probability (or transition probability), depending on the state of the system in each discrete time step Δ*t* = 1. The original definition ensures that the probabilities sum up to one so that one reaction (also reactions where no transition happens – when birth and death event happen within the same genotype) takes place in each time step. In the Moran process, a ‘payoff’ *π* is what a genotype gains from interactions with others. The interaction matrix is 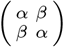, where *α* is the fitness gain for the parasite and the fitness loss for a host if the genotypes match, and *β* is the fitness gain or loss for mismatching pairs (here *α* = 1 and *β* = 0). For example, the probability of a *P*_1_ birth and thus a subsequent *P*_2_ death is proportional to *π*_*P*_1__ = *αh*_1_ + *βh*_2_, where *h*_1_ and *h*_2_, *p*_1_ and *p*_2_ are relative abundances. How much this effects the so-called ‘fitness’ *f* is controlled with the selection intensity *w_P_* (or *w_H_* for the host) so that *f*_*P*_1__ = 1 – *w_P_* + *w_P_ π*_*P*_1__. This per capita ‘fitness’ is then normalised by a dynamically changing population average 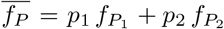 (this is the intraspecific competition step) and multiplied with the current abundance of the genotype. Thus 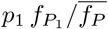 is then the birth probability of a genotype one parasite. The death probability is simply density dependent, thus the total probability of replacing a genotype-two (death) by a genotype-one (birth) parasite is 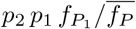. Since we are modelling two populations (host and parasite), we choose to update both populations simultaneously instead of sequentially, such that frequency changes of host genotypes and parasite genotypes can happen at once. The deterministic limit (population sizes *N_H_*, *N_P_* → ∞ and time steps Δ*t* → 0) of the Moran process in a single population is usually the differential equation of the replicator dynamics, however, in a two-population model the average fitness within each population is different and thus the *adjusted* replicator dynamics become the deterministic analogue [83, 65]. The adjusted replicator dynamics for host-parasite interactions have a globally attractive inner fixed point, in the symmetric matching alleles case this is the equal abundance of all genotypes.

**The discrete time Evo process (dtEvo)** [65, called pairwise comparison process or local update process in evolutionary game theory], is another birth death process, nearly equivalent to the Moran process but here competition is strictly local and pairwise, not normalised by a global average fitness. What is 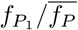 in the Moran process is here 0.5 + 0.5 (*f*_*P*_1__ – *f*_*P*_2__)/*max*(Δ*π_P_*). The ‘fitness’ of parasite 1 only depends on the difference in ‘fitness’ to parasite 2 which depends on the abundances of host genotypes (see equation for *f*_*P*_1__), but not, as when normalising with 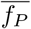, on the relative abundances of the parasite genotypes. Thus, in the pairwise comparison process, the antagonist influences globally (since there is no spatial structure), but within a species the competition is local. This results in the recovery of the replicator dynamics with neutral cycles in the deterministic limit.

**A Gillespie algorithm** [84] can be employed to simulate the above stochastic processes. In this case reaction rates (not probabilities) are calculated for each species and, using random numbers, the shortest waiting time for each reaction is determined under the assumption of exponential waiting times. The reaction with the shortest time takes place and time is updated accordingly. This makes time continuous and time steps unequal. In contrast to the discrete time models, the Gillespie algorithm only updates one species at a time. The now following processes are also implemented using a Gillespie algorithm.

**The EcoEvo process** uses independent reactions of host birth, parasite death and host-parasite interactions similar to the individual-based equivalent of the Lotka Volterra equations. This results in a microscopic process often believed to be a more natural approach because the reactions describe individual and independent events on the ‘microscopic’ level rather than population dynamics on the ‘macroscopic’ level. Host birth reactions are density dependent with constant rate *b_H_*, parasite death is density dependent with constant rate *d_P_* and a density-dependent interaction of matching host-parasite pairs can result in the death of the host or the birth of a parasite with constant rate λ. The population size has no restrictions in this case and freely follows the evolutionary dynamics. The deterministic analogue has a neutrally stable fixed point.

**The EcoEvo^+^ process** are like the EcoEvo independent reactions, but with additional competition in the host population. Density-dependent interactions of two host individuals, independent of the genotype, result in the death of one of the individuals with constant rate *μ*. This model, when reduced to the deterministic limit, is an antagonistic interaction model with logistic growth in the host (carrying capacity *K* = *b_H_*/*μ*) and an attractive inner fixed point.

**The Hybrid model** is a process with self-controlled population size. It is built on the EcoEvo model but with constrained birth and death rates adapted from the Evo model and dynamically varied to balance birth and death events on average. Thus, the population size is tightly controlled, yet it is not strictly constant. Building on the stochastic processes from evolutionary game theory above, one can set up a process that utilises the infection matrix for death events in the host and birth events in the parasite in explicit individual reactions. For example the dynamic reaction rate of a death event of a host genotype one is 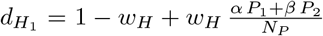. The birth rates for the host and the death rates for the parasite are then dynamically adjusted to equal the total rates of host death and parasite birth. The deterministic limit is the replicator dynamics with a neutrally stable fixed point, as in the pairwise comparison process.

## Authors’ contributions

Hinrich and Arne designed the research question. Hanna and Arne developed and adapted the models. Hanna conducted the analysis. All authors discussed and interpreted the results. Hanna wrote the initial draft. All authors revisited the manuscript critically and approved the final version.

## Acknowledgements

We want to thank Jens Christian Claussen and Andreas Rößler for detailed and enlightening discussions on methods, approximations and stochastic differential equations. We would like to thank Michael Raatz, Hye-Jin Park, Stefano Giaimo and Yuriy Pichugin for helpful comments on the manuscript.

