## supplemental file for "How long do Red Queen dynamics survive under genetic drift? A comparative analysis of evolutionary and eco-evolutionary models"

### Supplementary material

**S1 Additional results** **1**

**S2 Model descriptions** **3**

**S4 Deterministic properties** **11**

**S5 Approximate diffusion for a constant of motion** **11**

**S6 Exact sojourn times** **13**

**S7 Literature overview** **14**

### S1 Additional results

**Selection intensity:** Here, we analyse the impact of the selection intensity $w$ which modulates the pulling force in models with an attractive fixed point. For a more robust result we compare the simulations (lines in Figure S1) with sojourn times (section S6) calculated for the discrete time dtEvo$^+$ and dtEvo processes. Since only discrete time and discrete state processes can be represented by a transition matrix, we can only apply this approach to the discrete time constant population size processes. Although the Evo$^+$ process has an attracting fixed point, intuitively making extinction times longer than in the Evo process, for low population sizes and weak selection we see the opposite – fast extinction. Furthermore, while the extinction time increases exponentially with increasing intensity of selection in the Evo$^+$ process, the Evo extinction times stay comparably constant, which is not surprising considering the neutral stability. One interesting and unexpected result is that extinction time is lower for increasing selection intensity of one species while keeping the other constant. This occurs when one of the selection intensities is very low: some dark lines are decreasing, especially for example $w_H = 0.2$, and colours are reversed for low values of $w_P$.

**Dimensionality:** We next include more types to observe the exponential decline of diversity, here simply the number of genotypes $n_H$ and $n_P$ alive (Figure S2). The Evo$^+$ process simulated with a Gillespie algorithm is updated such that the interaction matrix is normalised depending on the number of strains (otherwise there is an imbalance between matching and non-matching pairs which results in change of selection strength). Oscillating selection can give an advantage to any type, but at different time points. A previously extinct parasite type ($P_2$) is reintroduced manually at time point 16, where the abundance of the corresponding host is especially high.

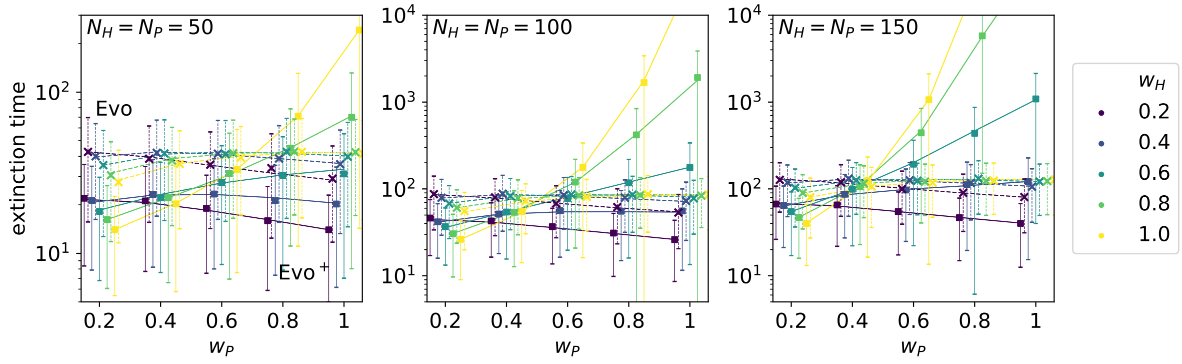

Figure S1: **Average extinction time for constant population sizes and different selection intensities.** The dtEvo$^+$ and dtEvo process are compared by the mean (■, ×) extinction times and the standard deviation from simulations with the exact sojourn times (—, - -) calculated analytically. The simulations start with equal abundance of both types $H_1(0) = H_2(0) = N_H/2$ and $P_1(0) = P_2(0) = N_P/2$. Parameters: $N_H = 250$, $w_H = w_P = 1$, $\alpha = 1$, $\beta = 0$. Note the log scale and different ranges on the y-axis.

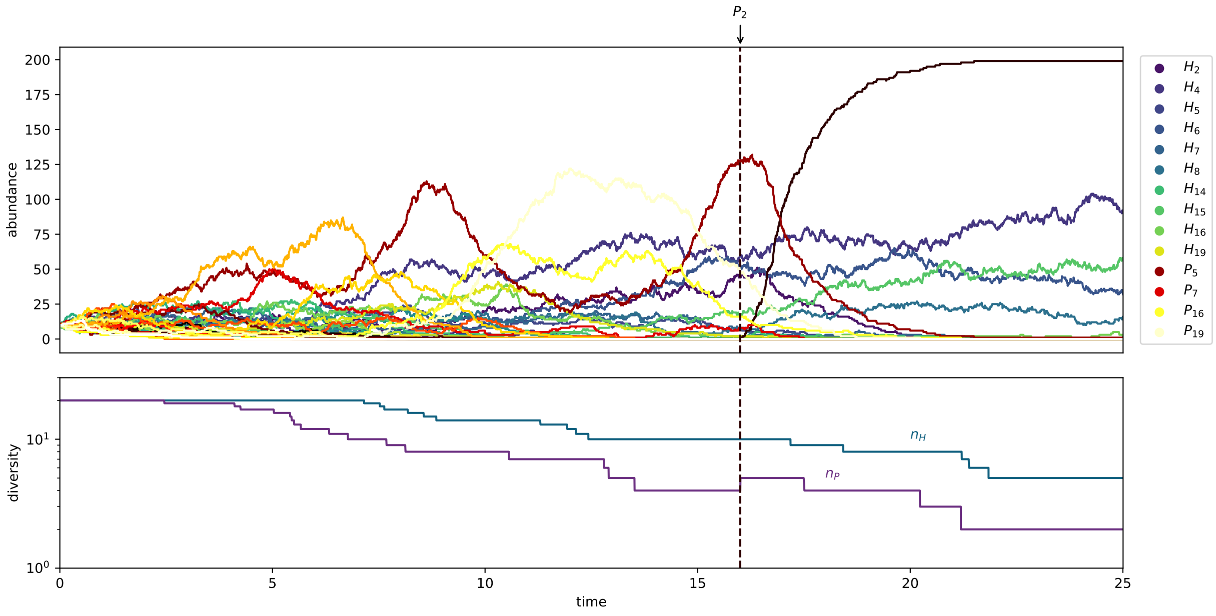

Figure S2: **Diversity decline** of subtypes of hosts and parasites. Example of an Evo$^+$ process implemented with a Gillespie algorithm. The simulations start with equal abundance of all 20 types $H_i(0) = N_H/20$ and $P_i(0) = N_P/20$. At time point $t = 16$ a well adapted but extinct $P_2 = 1$ is reintroduced manually, while $P_5$ is reduced by one individual to keep $N_P$ constant. Parameters: $N_H = 200$, $N_P = 200$, $w_H = w_P = 1$, $\alpha = 1$, $\beta = 0$. The legend shows host and parasite subtypes present at $t = 16$.

### S2 Model descriptions

Verbal model descriptions and some parameter explanations are available in the main text, here we show the mathematical details.

#### S2.1 The Evo$^+$ and Evo process

In the Evo$^+$ process (often Moran process in evolutionary game theory) and Evo process (often pairwise comparison or local update process) population size $N_H$ and $N_P$ is constant, so it is enough to focus on one type $H = H_1 = N_H - H_2$ and $P = P_1 = P_N - P_2$. Then the fitness of the second type has a negative influence on the first type and we can write fitness as $f_r^s$, with $s \in \{+, -\}$ indicating whether the fitness has a positive or negative influence on the focal type and $r \in \{h, p\}$ denoting the population. Fitness depends on the payoff taken from the game played between hosts and parasites with relative abundances $h := H/N_H$ and $p := P/N_P$. The infection matrices $M^p = \begin{pmatrix} \alpha & \beta \\ \beta & \alpha \end{pmatrix}$ and $M^h = \begin{pmatrix} \beta & \alpha \\ \alpha & \beta \end{pmatrix}$ result in payoffs

$$\begin{pmatrix} \pi_p^+ \\ \pi_p^- \end{pmatrix} = M^p \begin{pmatrix} h \\ 1-h \end{pmatrix} = \begin{pmatrix} \alpha h+\beta(1-h) \\ \beta h+\alpha(1-h) \end{pmatrix}$$

and

$$\begin{pmatrix} \pi_h^+ \\ \pi_h^- \end{pmatrix} = M^h \begin{pmatrix} p \\ 1-p \end{pmatrix} = \begin{pmatrix} \beta p+\alpha(1-p) \\ \alpha p+\beta(1-p) \end{pmatrix}.$$

The strength of the influence of this payoff on fitness is modulated by the selection intensity $w_r \in [0, 1]$. Here we use a linear dependence

$$f_r^s = 1 - w_r + w_r \pi_r^s. \tag{1}$$

In an Evo$^+$ process the local probabilities depend on the fitness of a type divided by an average fitness (fitness of subtypes weighed with the relative abundance) of the species.

$$\Phi_r^s = \frac{f_r^s}{\langle f_r \rangle} = \frac{f_r^s}{r f_r^+ + (1-r) f_r^-}. \tag{2}$$

For the Evo process, the local transition probabilities are influenced by the fitness difference between the two types scaled with the maximal possible payoff difference

$$\Phi_r^s = \frac{1}{2} + \frac{1}{2} \frac{f_r^s - f_r^{\neg s}}{\Delta \pi_{r_{max}}}, \tag{3}$$

where $\neg s$ is the other type (chosen for death). We often use $\alpha = 1$ and $\beta = 0$, then $\Delta\pi_{r_{max}} = 1$, else $\Delta\pi_{r_{max}} = \alpha - \beta$. The local rates $\Phi_r^s$ are the specific rates for the birth-death reactions. Multiplied with the relative abundance of the reactants, we get the global transition probabilities $T_r^s = r\,(1-r)\,\Phi_r^s$.

In the discrete time processes host and parasite populations are updated simultaneously and all transition probabilities sum up to one. For each population, there must therefore be a transition probability for increasing, decreasing or not changing the number of the focal type. The respective probabilities are $T_r^+$, $T_r^-$ and $T_r^0 = 1 - T_r^+ - T_r^-$. For the two interacting populations this yields nine probabilities, Tab. S1.

In the computer simulation, in each time step $\Delta t = 1$, a random number is compared with the cumulative probabilities and the resulting transition is carried out by adjusting the population accordingly and updating time to $t + 1$.

#### S2.2 Gillespie algorithm

In a Gillespie algorithm we assume independent interactions with one or two reactants that react with a certain rate, which is the product of the reactant concentrations multiplied with a reaction constant (not

| Transition in (h,p)-space | number $l$ | $\Delta h$ | $\Delta p$ | Probability $\mathcal{T}_l$ |
|---|---|---|---|---|
| $\bullet$ | 1 | 0 | 0 | $T_h^0\, T_p^0$ |
| $\rightarrow$ | 2 | $\frac{1}{N_H}$ | 0 | $T_h^+\, T_p^0$ |
| $\uparrow$ | 3 | 0 | $\frac{1}{N_P}$ | $T_h^0\, T_p^+$ |
| $\leftarrow$ | 4 | $-\frac{1}{N_H}$ | 0 | $T_h^-\, T_p^0$ |
| $\downarrow$ | 5 | 0 | $-\frac{1}{N_P}$ | $T_h^0\, T_p^-$ |
| $\nearrow$ | 6 | $\frac{1}{N_H}$ | $\frac{1}{N_P}$ | $T_h^+\, T_p^+$ |
| $\swarrow$ | 7 | $-\frac{1}{N_H}$ | $-\frac{1}{N_P}$ | $T_h^-\, T_p^-$ |
| $\searrow$ | 8 | $\frac{1}{N_H}$ | $-\frac{1}{N_P}$ | $T_h^+\, T_p^-$ |
| $\nwarrow$ | 9 | $-\frac{1}{N_H}$ | $\frac{1}{N_P}$ | $T_h^-\, T_p^+$ |

Table S1: **Transition probabilities** $\mathcal{T}_l(h,p)$

necessarily a constant here). If reactions happen with waiting times that follow an exponential distribution we can calculate the time of the next reaction, for each reaction with mass-action rate $\phi$ and the help of a random number $R \in [0,1]$,

$$\Delta t = \frac{1}{\phi} \log \frac{1}{R}. \tag{4}$$

We then choose the reaction that happens first and update population and time.

For reactions with one reactant, the reaction rates are multiplied with the absolute abundance (e.g. $\phi = H_1 b$ for the first reaction in the EcoEvo$^+$ process), but when two reactants are necessary, the probability of that reaction would be amplified. The reaction is therefore scaled. In the Evo$^+$/Evo process pair reactions are scaled with $N_H$ ($N_P$) for the first two (last two) reactions (see Table S2), so that $\phi = H_1 \frac{H_2}{N_H} \Phi_H^+$, etc. In the EcoEvo$^+$ and EcoEvo process reactions with two reactants are scaled with the carrying capacity $K$ ($\phi = H_1 \frac{H_2}{K} \mu$ for the third reaction). The last case is very close to the usual approach in chemistry, where reactions rates with more than one reactant are scaled with the system volume.

#### S2.3 EcoEvo$^+$ and EcoEvo process

For the completely independent EcoEvo models reactions with two reactants are scaled with an extrinsic carrying capacity $K$. These are reactions where host and parasite interact with rate $\lambda = \frac{\lambda_0}{K}$ or where hosts are in competition with each other $\mu = \frac{b_h}{K}$ (or $\mu = 0$ in the IR model). The choice of $\mu$ becomes apparent in the deterministic limit, where the carrying capacity in the logistic term is $K = \frac{b_h}{\mu}$ (see Tab. S3). Note that the carrying capacity does not equal the population size of the hosts when parasites are present. The initial conditions are chosen to be similar with the game theory processes by varying the value of the fixed point through $b_h$.

#### S2.4 Hybrid process

Like the Evo$^+$ an Evo process the Hybrid model takes its interaction rates via a game theory approach from the payoff matrix. In the Evo$^+$ and Evo process the payoff matrix for the host is adjusted to realise positive fitness. Here, the set-up is closer to typical host-parasite modelling by using only one payoff matrix for positive rates $b_{P_1}$, $b_{P_2}$ for the parasite and negative rates $d_{H_1}$, $d_{H_2}$ for the host. With $M = \begin{pmatrix} \alpha & \beta \\ \beta & \alpha \end{pmatrix}$ we obtain $b_{P_1} = 1 - w_P + w_P \frac{H_1\alpha + H_2\beta}{H_1+H_2}$, $b_{P_2} = 1 - w_P + w_P \frac{H_1\beta + H_2\alpha}{H_1+H_2}$, $d_{H_1} = 1 - w_H + w_H \frac{P_1\alpha + P_2\beta}{P_1+P_2}$ and

$d_{H_2} = 1 - w_H + w_H \frac{P_1\beta + P_2\alpha}{P_1+P_2}$. When $w_H = w_P = 1$ the model is equivalent to the independent reactions with two reactants, however, scaled with different quantities $H_1 + H_2$ or $P_1 + P_2$, which are dynamic. This model is in between the EcoEvo and the Evo$^+$/Evo processes.

| Model | reactions | time | population size | fixed point | dimension |
|---|---|---|---|---|---|
| discrete time Evo$^+$<br><br>(dtEvo$^+$) | $H_1 + H_2 \xrightarrow{T_P^0 \Phi_H^+} 2H_1$<br>$H_1 + H_2 \xrightarrow{T_P^0 \Phi_H^-} 2H_2$<br>$P_1 + P_2 \xrightarrow{T_H^0 \Phi_P^+} 2P_1$<br>$P_1 + P_2 \xrightarrow{T_H^0 \Phi_P^-} 2P_2$<br>$H_1 + H_2 + P_1 + P_2 \xrightarrow{\Phi_H^+ \Phi_P^+} 2H_1 + 2P_1$<br>$H_1 + H_2 + P_1 + P_2 \xrightarrow{\Phi_H^+ \Phi_P^-} 2H_1 + 2P_2$<br>$H_1 + H_2 + P_1 + P_2 \xrightarrow{\Phi_H^- \Phi_P^+} 2H_2 + 2P_1$<br>$H_1 + H_2 + P_1 + P_2 \xrightarrow{\Phi_H^- \Phi_P^-} 2H_2 + 2P_2$<br>no reaction with probability $T_H^0 T_P^0$ | discrete | constant | attractive | 2D |
| Evo$^+$ | $H_1 + H_2 \xrightarrow{\Phi_H^+} 2H_1$<br>$H_1 + H_2 \xrightarrow{\Phi_H^-} 2H_2$<br>$P_1 + P_2 \xrightarrow{\Phi_P^+} 2P_1$<br>$P_1 + P_2 \xrightarrow{\Phi_P^-} 2P_2$ | continuous | constant | attractive | 2D |
| discrete time Evo<br><br>dtEvo | like discrete time Evo$^+$, but different equations for $\Phi$ and $T$ | discrete | constant | neutral | 2D |
| Evo | like Gillespie Evo$^+$, with different $\Phi$ and $T$ | continuous | constant | neutral | 2D |
| Hybrid | $H_1 \xrightarrow{(H_1 d_{H_1} + H_2 d_{H_2})/(H_1+H_2)} 2H_1$<br>$H_2 \xrightarrow{(H_1 d_{H_1} + H_2 d_{H_2})/(H_1+H_2)} 2H_2$<br>$\left.\begin{array}{l} H_1 \xrightarrow{d_{H_1}} \emptyset \\ H_2 \xrightarrow{d_{H_2}} \emptyset \\ P_1 \xrightarrow{b_{P_1}} 2P_1 \\ P_2 \xrightarrow{b_{P_2}} 2P_2 \end{array}\right\}$<br>$P_1 \xrightarrow{(P_1 b_{P_1} + P_2 b_{P_2})/(P_1+P_2)} \emptyset$<br>$P_2 \xrightarrow{(P_1 b_{P_1} + P_2 b_{P_2})/(P_1+P_2)} \emptyset$<br>death host and birth parasite when $w_H = w_P = 1$ :<br>$H_1 + P_1 \xrightarrow{\alpha} P_1$<br>$H_1 + P_2 \xrightarrow{\beta} P_2$<br>$H_2 + P_1 \xrightarrow{\beta} P_1$<br>$H_2 + P_2 \xrightarrow{\alpha} P_2$<br>$P_1 + H_1 \xrightarrow{\alpha} 2P_1 + H_1$<br>$P_1 + H_2 \xrightarrow{\beta} 2P_1 + H_2$<br>$P_2 + H_1 \xrightarrow{\beta} 2P_2 + H_1$<br>$P_2 + H_2 \xrightarrow{\alpha} 2P_2 + H_2$ | continuous | nearly constant | neutral | 4D |
| EcoEvo$^+$ | $H_1 \xrightarrow{b} 2H_1$<br>$H_2 \xrightarrow{b} 2H_2$<br>$H_1 + H_2 \xrightarrow{\mu} H_1$<br>$H_1 + H_1 \xrightarrow{\mu} H_1$<br>$H_1 + H_2 \xrightarrow{\mu} H_2$<br>$H_2 + H_2 \xrightarrow{\mu} H_2$<br>$P_1 + H_1 \xrightarrow{\lambda} P_1$<br>$P_1 + H_1 \xrightarrow{\lambda} 2P_1 + H_1$<br>$P_2 + H_2 \xrightarrow{\lambda} P_2$<br>$P_2 + H_2 \xrightarrow{\lambda} 2P_2 + H_2$<br>$P_1 \xrightarrow{d} \emptyset$<br>$P_2 \xrightarrow{d} \emptyset$ | continuous | constrained | attractive | 4D |
| EcoEvo | like EcoEvo$^+$ but with $\mu = 0$ | continuous | unconstrained | neutral | 4D or 2×2D |

Table S2: **Model overview.** Model names and their main properties. See main text for explanation.

### S3 Stochastic differential equation and deterministic limit

#### S3.1 Master equation

In the limit of $N_H, N_P \to \infty$ and $\Delta t \to 0$ the stochastic process is described by a Fokker Planck equation with a selection term (the deterministic ordinary differential equation, often called drift in physics) and a diffusion term (stochastic term often called genetic drift in biology). The Fokker Planck equation can also be converted into a stochastic differential equation. As an example we explain this procedure for the discrete time Evo$^+$ or Evo process.

From the transition probabilities above, one can write down the master equation

$$\begin{aligned} P(t+\Delta t, h, p) =& P(t, h, p) \\ &+ \Delta t \sum_{l=2}^{9} P(t, h-\Delta h, p-\Delta p)\, \mathcal{T}_l(h-\Delta h, p-\Delta p) \\ &- \Delta t\, P(t, h, p) \sum_{l=2}^{9} \mathcal{T}_l(h, p) \end{aligned} \tag{5}$$

The probability $P(h, p, t+\Delta t)$ of being at the specific state $(h, p)$ at time $t+\Delta t$ is the probability of being in that state plus the probability of being in a state close by at time $t$ multiplied with the probability of transitioning to the new state $(h, p)$ minus the probability of leaving that state. See Tab. S1.

#### S3.2 Fokker-Planck equation

We can now use the shift operator $E_{\Delta x}$

$$\begin{aligned} f(x+\Delta x) &= E_{\Delta x} f(x) = e^{\Delta x \frac{d}{dx}} f(x) \\ &= \left(1 + \Delta x \frac{d}{dx} + \Delta x^2 \frac{1}{2!} \frac{d^2}{dx^2} + ...\right) f(x) \end{aligned} \tag{6}$$

on the left hand side and on the right hand side of the master equation. Then approximating

$$P(t+\Delta t, h, p) - P(t, h, p) = \frac{d}{dt} \Delta t P(t, h, p) \tag{7}$$

to first order and

$$\begin{aligned} P(t,&h-\Delta h, p-\Delta p) \mathcal{T}_l(h-\Delta h, p-\Delta p) \\ &= \left(1 - \Delta h \frac{d}{dh} + \Delta h \frac{d^2}{dh^2}\right) \left(1 - \Delta p \frac{d}{dp} + \Delta p \frac{d^2}{dp^2}\right) P(t, h, p) \mathcal{T}_l(h, p) \end{aligned} \tag{8}$$

to second order ($\frac{1}{N^2}$, see (van Kampen, 1997; Risken, 1996; Gardiner, 1985)) we get

$$
\begin{aligned}
\frac{d}{dt}P(t,h,p) = & \\
&+\left(-\frac{1}{N_H}\frac{\partial}{\partial h} + \frac{1}{2N_H^2}\frac{\partial^2}{\partial h^2}\right) P(t,h,p)\mathcal{T}_2(h,p) \\
&+\left(-\frac{1}{N_P}\frac{\partial}{\partial p} + \frac{1}{2N_P^2}\frac{\partial^2}{\partial p^2}\right) P(t,h,p)\mathcal{T}_3(h,p) \\
&+\left(+\frac{1}{N_H}\frac{\partial}{\partial h} + \frac{1}{2N_H^2}\frac{\partial^2}{\partial h^2}\right) P(t,h,p)\mathcal{T}_4(h,p) \\
&+\left(+\frac{1}{N_P}\frac{\partial}{\partial p} + \frac{1}{2N_P^2}\frac{\partial^2}{\partial p^2}\right) P(t,h,p)\mathcal{T}_5(h,p) \\
&+\left(-\frac{1}{N_H}\frac{\partial}{\partial h} - \frac{1}{N_P}\frac{\partial}{\partial p} + \frac{1}{2N_H^2}\frac{\partial^2}{\partial h^2} + \frac{1}{N_H N_P}\frac{\partial}{\partial h}\frac{\partial}{\partial p} + \frac{1}{2N_P^2}\frac{\partial^2}{\partial p^2}\right) P(t,h,p)\mathcal{T}_6(h,p) \\
&+\left(+\frac{1}{N_H}\frac{\partial}{\partial h} + \frac{1}{N_P}\frac{\partial}{\partial p} + \frac{1}{2N_H^2}\frac{\partial^2}{\partial h^2} + \frac{1}{N_H N_P}\frac{\partial}{\partial h}\frac{\partial}{\partial p} + \frac{1}{2N_P^2}\frac{\partial^2}{\partial p^2}\right) P(t,h,p)\mathcal{T}_7(h,p) \\
&+\left(-\frac{1}{N_H}\frac{\partial}{\partial h} + \frac{1}{N_P}\frac{\partial}{\partial p} + \frac{1}{2N_H^2}\frac{\partial^2}{\partial h^2} - \frac{1}{N_H N_P}\frac{\partial}{\partial h}\frac{\partial}{\partial p} + \frac{1}{2N_P^2}\frac{\partial^2}{\partial p^2}\right) P(t,h,p)\mathcal{T}_8(h,p) \\
&+\left(+\frac{1}{N_H}\frac{\partial}{\partial h} - \frac{1}{N_P}\frac{\partial}{\partial p} + \frac{1}{2N_H^2}\frac{\partial^2}{\partial h^2} - \frac{1}{N_H N_P}\frac{\partial}{\partial h}\frac{\partial}{\partial p} + \frac{1}{2N_P^2}\frac{\partial^2}{\partial p^2}\right) P(t,h,p)\mathcal{T}_9(h,p).
\end{aligned}
$$

The resulting Fokker-Planck equation emerges when we collect similar terms

$$
\begin{aligned}
\frac{\partial}{\partial t}P(t,h,p) &= \left(-\frac{\partial}{\partial h}\,a_1 - \frac{\partial}{\partial p}\,a_2\right) P(t,h,p) \\
&\quad \frac{1}{2}\left(+\frac{\partial^2}{\partial h^2}\,d_1 + \frac{\partial^2}{\partial p^2}\,d_2 + \frac{\partial}{\partial h}\frac{\partial}{\partial p}\,d_{1,2} + \frac{\partial}{\partial p}\frac{\partial}{\partial h}\,d_{2,1}\right) P(t,h,p).
\end{aligned}
\tag{9}
$$

The drift vector $\boldsymbol{a} = (a_1, a_2)$ and the diffusion matrix $\boldsymbol{D} = \begin{pmatrix} d_1 & d_{1,2} \\ d_{2,1} & d_2 \end{pmatrix}$ are calculated as follows, where we have dropped the arguments $(h,p)$ for readability.

$$
\begin{aligned}
a_1 = \dot{h} &= \frac{1}{N_H}\left(\mathcal{T}_2 + \mathcal{T}_6 + \mathcal{T}_8 - \mathcal{T}_4 - \mathcal{T}_7 - \mathcal{T}_9\right) = \frac{1}{N_H}(T_H^+ - T_H^-) \\
&= \frac{h(1-h)}{N_H}(\Phi_H^+ - \Phi_H^-) \\
a_2 = \dot{p} &= \frac{1}{N_P}\left(\mathcal{T}_3 + \mathcal{T}_6 + \mathcal{T}_9 - \mathcal{T}_5 - \mathcal{T}_7 - \mathcal{T}_8\right) = \frac{p(1-p)}{N_P}(\Phi_P^+ - \Phi_P^-) \\
d_1 &= \frac{1}{N_H^2}\left(\mathcal{T}_2 + \mathcal{T}_4 + \mathcal{T}_6 + \mathcal{T}_7 + \mathcal{T}_8 + \mathcal{T}_9\right) = \frac{h(1-h)}{N_H^2}(\Phi_H^+ + \Phi_H^-) \\
d_2 &= \frac{1}{N_P^2}\left(\mathcal{T}_3 + \mathcal{T}_5 + \mathcal{T}_6 + \mathcal{T}_7 + \mathcal{T}_8 + \mathcal{T}_9\right) = \frac{p(1-p)}{N_P^2}(\Phi_P^+ + \Phi_P^-) \\
d_{1,2} = d_{2,1} &= \frac{1}{N_H N_P}\left(\mathcal{T}_6 + \mathcal{T}_7 - \mathcal{T}_8 - \mathcal{T}_9\right) \\
&= \frac{h(1-h)p(1-p)}{N_H N_P}\left(\Phi_H^+ + \Phi_H^-\right)\left(\Phi_P^+ + \Phi_P^-\right).
\end{aligned}
$$

We have summarised the results for the infection matrices $M^p = \begin{pmatrix} \alpha & \beta \\ \beta & \alpha \end{pmatrix}$ and $M^h = \begin{pmatrix} \beta & \alpha \\ \alpha & \beta \end{pmatrix}$ in Tab. S3. For the Evo$^+$ process $\Phi_H^+ - \Phi_H^- = \frac{w_H(\alpha-\beta)(1-2p)}{1-w_H+w_H\langle\pi_H\rangle}$, $\Phi_H^+ + \Phi_H^- = \frac{2-2w_H+w_H(\alpha+\beta)}{1-w_H+w_H\langle\pi_H\rangle}$, $\Phi_P^+ - \Phi_P^- = \frac{w_P(\alpha-\beta)(2h-1)}{1-w_P+w_P\langle\pi_P\rangle}$ and $\Phi_P^+ + \Phi_P^- = \frac{2-2w_P+w_P(\alpha+\beta)}{1-w_P+w_P\langle\pi_P\rangle}$. And for the Evo process $\Phi_H^+ - \Phi_H^- = w_H(1-2p)$, $\Phi_H^+ + \Phi_H^- = 1$,

$\Phi_P^+ - \Phi_P^- = w_P(2h-1)$ and $\Phi_P^+ + \Phi_P^- = 1$. We have normalised the reaction rates in the discrete time processes so that all probabilities sum up to one. The other processes' reaction rates are on the order of $N_H$ or $N_P$ (also see appendix section S2.2), for the continuous time processes the Fokker Planck equation is therefore faster. Furthermore, some of the processes are four dimensional with drift vector $\boldsymbol{a} = (a_1, a_2, a_3, a_4)$ for $H_1$, $H_2$, $P_1$, $P_2$ and diffusion matrix $\boldsymbol{D} = \begin{pmatrix} d_1 & 0 & 0 & 0 \\ 0 & d_2 & 0 & 0 \\ 0 & 0 & d_3 & 0 \\ 0 & 0 & 0 & d_4 \end{pmatrix}$. Here, all correlative noise (non-diagonal entries) is zero because all the reactions only change one reactant at a time. Note that without diffusion, the Hybrid process is equivalent to the scaled two-dimensional replicator dynamics because $a_1 + a_2 = \dot{H}_1 + \dot{H}_2 = 0$ which implies $H_1 + H_2 = N_H$.

#### S3.3 Stochastic differential equation

From the Fokker Planck equation a stochastic differential equation (SDE) can be constructed. The SDE is a differential equation with a deterministic part, which has to be integrated with respect to time and a second term which is integrated with the Wiener process (continuous time random walk/Brownian motion). While the Fokker Planck equation describes the probabilities for certain states in time, and thus a probability distribution, the SDE accounts for the change of a random variable. Integrated with respect to one particular realisation of a random walk (Wiener process), it will give one particular realisation of the stochastic process.

To derive a non-unique SDE from the Fokker Planck equation we take the "square root" of the matrix $D = \begin{pmatrix} d_1 & d_{1,2} \\ d_{2,1} & d_2 \end{pmatrix} = B^T B$ with $d_{1,2} = d_{2,1}$. A Cholesky decomposition can achieve this,

$$B = \begin{pmatrix} \sqrt{d_1} & \frac{d_{1,2}}{\sqrt{d_1}} \\ 0 & \sqrt{d_2 - \frac{d_{1,2} c\left(\frac{d_{1,2}}{\sqrt{d_1}}\right)}{\sqrt{d_1}}} \end{pmatrix} \tag{10}$$

where $c(x)$ is the complex conjugate of $x$. This is only important if $d_1$ can become negative which is not the case for our purposes. With the drift term $a(X_t)$ diffusion matrix $B = B(X_t)$ we set up the stochastic differential equation in the Itô sense

$$dX_t = a(X_t)\, dt + B(X_t)\, dW_t \tag{11}$$

where the stochastic variable $X_t = \binom{h}{p}$ describes the relative abundance of the first host and first parasite, and $W$ is the Wiener process. The terms $a_1$ and $a_2$ are equivalent to the differential equation (deterministic terms in Table S3) and describe the $N \to \infty$ limit of the stochastic process, which is e.g. the replicator equation for the Evo process and the adjusted replicator dynamics for the Evo$^+$ process.

The Fokker Planck equation is a deterministic partial differential equation, which can be solved in simple cases. Although this is not the case here, we can learn from the matrix $D$ how the variance in the distribution is affected by the parameters. Again, although all seven models stem from the same biological mechanisms, the results are quite different. The Evo$^+$, Evo, EcoEvo$^+$ and EcoEvo processes have only diagonal elements, therefore the random term is not correlated across species (2D models) or even types (4D models).

Stochastic differential equations can be numerically integrated in a robust way (Rößler, 2010; Kloeden and Platen, 1992) using random numbers for the Wiener process. The advantage of an SDE with respect to a stochastic simulation (via Gillespie's algorithm) is the independence of run time on the population size $N$. In a stochastic simulation each individual in the population is simulated. An SDE, like an ODE is integrated stepwise, thus the limiting factor here is the time step used. For small population numbers as displayed in the figures of the main text both algorithms are comparable in speed. As population size increases stochastic simulations become infeasible, yet SDE integration does not increase the runtime.

| Model | deterministic terms | stochastic terms |
|---|---|---|
| dtEvo$^+$ | $\dot{h} = \frac{h(1-h)}{N_H} \frac{w_H(\alpha-\beta)(1-2p)}{1-w_H+w_H\langle\pi_H\rangle}$<br>$\dot{p} = \frac{p(1-p)}{N_P} \frac{w_P(\alpha-\beta)(2h-1)}{1-w_P+w_P\langle\pi_P\rangle}$ | $d_1 = \frac{h(1-h)}{N_H^2} \frac{2-2w_H+w_H(\alpha+\beta)}{1-w_H+w_H\langle\pi_H\rangle}$<br>$d_2 = \frac{p(1-p)}{N_P^2} \frac{2-2w_P+w_P(\alpha+\beta)}{1-w_P+w_P\langle\pi_P\rangle}$<br>$d_{1,2} = \frac{h(1-h)p(1-p)}{N_H N_P}$<br>$\frac{2-2w_H+w_H(\alpha+\beta)}{1-w_H+w_H\langle\pi_H\rangle} \frac{2-2w_P+w_P(\alpha+\beta)}{1-w_P+w_P\langle\pi_P\rangle}$ |
| Evo$^+$ | $\dot{h} = h(1-h)\frac{w_H(\alpha-\beta)(1-2p)}{1-w_H+w_H\langle\pi_H\rangle}$<br>$\dot{p} = p(1-p)\frac{w_P(\alpha-\beta)(2h-1)}{1-w_P+w_P\langle\pi_P\rangle}$ | $d_1 = \frac{h(1-h)}{N_H} \frac{2-2w_H+w_H(\alpha+\beta)}{1-w_H+w_H\langle\pi_H\rangle}$<br>$d_2 = \frac{p(1-p)}{N_P} \frac{2-2w_P+w_P(\alpha+\beta)}{1-w_P+w_P\langle\pi_P\rangle}$<br>$d_{1,2} = 0$ |
| dtEvo | $\dot{h} = \frac{h(1-h)}{N_H} w_H(1-2p)$<br>$\dot{p} = \frac{p(1-p)}{N_P} w_P(2h-1)$ | $d_1 = \frac{h(1-h)}{N_H^2}$<br>$d_2 = \frac{p(1-p)}{N_P^2}$<br>$d_{1,2} = \frac{h(1-h)p(1-p)}{N_H N_P}$ |
| Evo | $\dot{h} = h(1-h)w_H(1-2p)$<br>$\dot{p} = p(1-p)w_P(2h-1)$ | $d_1 = \frac{h(1-h)}{N_H}$<br>$d_2 = \frac{p(1-p)}{N_P}$<br>$d_{1,2} = 0$ |
| Hybrid | $\dot{H}_1 = \frac{H_1 H_2 w_H(\alpha-\beta)(P_2-P_1)}{(H_1+H_2)(P_1+P_2)}$<br>$\dot{H}_2 = \frac{H_1 H_2 w_H(\alpha-\beta)(P_1-P_2)}{(H_1+H_2)(P_1+P_2)}$<br>$\dot{P}_1 = \frac{P_1 P_2 w_P(\alpha-\beta)(H_1-H_2)}{(H_1+H_2)(P_1+P_2)}$<br>$\dot{P}_2 = \frac{P_1 P_2 w_P(\alpha-\beta)(H_2-H_1)}{(H_1+H_2)(P_1+P_2)}$ | $d_1 = 2H_1(1 - w_H + w_H(\alpha(P_1 H_1 +$<br>$H_2 \frac{P_1+P_2}{2}) + \beta(P_2 H_1 + H_2 \frac{P_1+P_2}{2})))$<br>$d_2 = 2H_2(1 - w_H + w_H(\alpha(P_2 H_2 +$<br>$H_1 \frac{P_1+P_2}{2}) + \beta(P_1 H_2 + H_1 \frac{P_1+P_2}{2})))$<br>$d_3 = 2P_1(1 - w_P + w_P(\alpha(P_1 H_1 +$<br>$P_2 \frac{H_1+H_2}{2}) + \beta(H_2 P_1 + P_2 \frac{H_1+H_2}{2})))$<br>$d_4 = 2P_2(1 - w_P + w_P(\alpha(P_2 H_2 +$<br>$P_1 \frac{H_1+H_2}{2}) + \beta(H_1 P_2 + P_1 \frac{H_1+H_2}{2})))$ |
| EcoEvo$^+$ | $\dot{H}_1 = b_h H_1(1 - \frac{\mu}{b_h}(H_1 + H_2))$<br>$-\lambda P_1 H_1$<br>$\dot{H}_2 = b_h H_2(1 - \frac{\mu}{b_h}(H_1 + H_2))$<br>$-\lambda P_2 H_2$<br>$\dot{P}_1 = P_1(\lambda H_1 - d_p)$<br>$\dot{P}_2 = P_2(\lambda H_2 - d_p)$ | $d_1 = b_h H_1(1 + \frac{\mu}{b_h}(H_1 + H_2)) + \lambda P_1 H_1$<br>$d_2 = b_h H_2(1 + \frac{\mu}{b_h}(H_1 + H_2)) + \lambda P_2 H_2$<br>$d_3 = \lambda P_1 H_1 + d_p P_1$<br>$d_4 = \lambda P_2 H_2 + d_p P_2$ |
| EcoEvo | $\dot{H}_1 = b_h H_1 - \lambda P_1 H_1$<br>$\dot{H}_2 = b_h H_2 - \lambda P_2 H_2$<br>$\dot{P}_1 = P_1(\lambda H_1 - d_p)$<br>$\dot{P}_2 = P_2(\lambda H_2 - d_p)$ | $d_1 = b_h H_1 + \lambda P_1 H_1$<br>$d_2 = b_h H_2 + \lambda P_2 H_2$<br>$d_3 = \lambda P_1 H_1 + d_p P_1$<br>$d_4 = \lambda P_2 H_2 + d_p P_2$ |

Table S3: **Deterministic and stochastic terms** for the time evolution of relative abundances $h = H_1/N_H$ or absolute abundances $H_1$, etc. for all models. Deterministic equations are the drift terms in the Fokker Planck equation. Stochastic terms are the diffusion terms in the Fokker Planck equation. See main text for explanation.

### S4 Deterministic properties

**Fixed points** occur when the dynamics have come to a halt and there is no more change in (relative) abundances. In the two-dimensional deterministic models (dtEvo$^+$, Evo$^+$, dtEvo, Evo and Hybrid - if normalised), which can be described by the replicator dynamics or the adjusted replicator dynamics, there are trivial points, where one subtype is extinct: ($h = 0$ or $h = 1$) and ($p = 0$ or $p = 1$). The less trivial fixed point is the inner coexistence fixed point with $h^* = p^* = 0.5$ in the symmetric case where there is no distinction between subtypes except the specificity to a matching host/parasite. The four-dimensional deterministic models (EcoEvo$^+$ and EcoEvo) have coexistence fixed points $H_i^* = \frac{d_p}{\lambda} = \frac{d_p\, K}{\lambda_0}$ and $P_i^* = \frac{b_h}{\lambda}\left(1 - \frac{\mu}{b_h}(H_1^* + H_2^*)\right)$, where $P_i^* = \frac{b_h}{\lambda}$ when $\mu = 0$, there is no logistic growth.
**A linear stability analysis** is carried out to learn more about the properties of the fixed point. The Jacobian at this inner fixed point $J(h^*, p^*) = \left.\begin{pmatrix} \frac{d\dot{h}}{dh} & \frac{d\dot{h}}{dp} \\ \frac{d\dot{p}}{dh} & \frac{d\dot{p}}{dp} \end{pmatrix}\right|_{h=h^*, p=p^*}$ is calculated to reveal the characteristic function $\det\left(J(h^*, p^*) - \lambda I\right) = 0$. If the real parts of all Eigenvalues are negative, the fixed point is attractive and thus (in the terms of evolutionary game theory) an evolutionary stable strategy ESS. If at least one Eigenvalue has a positive real part, the state is a repellor (unstable). And if all real parts are zero, it is not enough to conduct a linear stability analysis. For the replicator dynamics and the adjusted replicator dynamics we find only imaginary Eigenvalues where the real part is zero, indicating neutral stability. This can be addressed by more sophisticated methods.

A constant of motion (Hofbauer, 1996; Koth and Sigmund, 1987) for the replicator dynamics reveals that the dynamics are volume preserving. The fixed point is neutrally stable. All trajectories follow orbits as shown in Fig. S3.
Also, from Hofbauer and Sigmund (1998, chapter 11) we know that in our example with $M^p = \begin{pmatrix} \alpha & \beta \\ \beta & \alpha \end{pmatrix}$ and $M^h = \begin{pmatrix} \beta & \alpha \\ \alpha & \beta \end{pmatrix}$ we have a *c-zero-sum* game. Then the fixed point is globally asymptotically stable (attractive) in the adjusted replicator dynamics.
A simple stability analysis shows that the EcoEvo$^+$ is attractive for many parameters but neutral when $\mu = 0$ in the EcoEvo interactions with no constraint on population size.

### S5 Approximate diffusion for a constant of motion

For the neutrally stable models that lead to replicator dynamics (dtEvo, Evo, Hybrid) and to the original independent reactions dynamics (EcoEvo) one can set up a constant of motion $H$. Here we use the replicator dynamics as an example.

A constant of motion is a function of the state variables in the system that does not change in time $\frac{dH}{dt} = 0$ when a trajectory is not perturbed by chance. Intuitively one can think of a system with constant energy where the dynamics are confined to the constant energy contour lines. For the two-dimensional replicator dynamics and a matching allele infection matrix $M^p = \begin{pmatrix} 1 & 0 \\ 0 & 1 \end{pmatrix}$ and $M^h = \begin{pmatrix} 0 & 1 \\ 1 & 0 \end{pmatrix}$ the constant of motion is

$$H(h, p) = 16\, h\, p\, (1 - h)\, (1 - p) \tag{12}$$

$h, p \in [0, 1]$ are the relative abundances of host type one and parasite type one ($1 - h$ and $1 - p$ are the relative abundances of type two) and the constant of motion is scaled so that it has the value one when both types are equally abundant ($h = p = 1/2$) and the value zero when any type is extinct (Claussen, 2016). The deterministic trajectory would start with a certain value of $H$, depending on the initial condition, and would follow that line indefinitely (see Fig. S3).

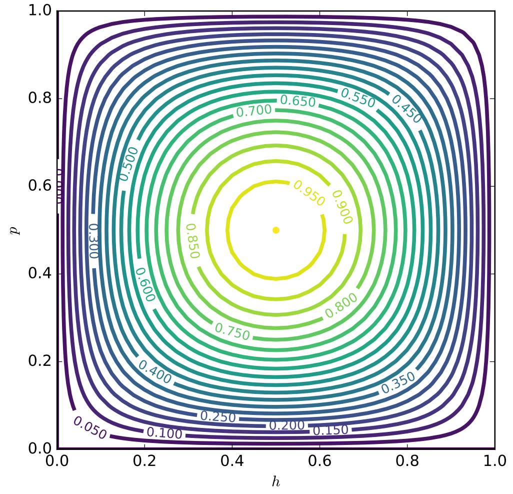

Figure S3: **Constant of motion for replicator dynamics.** Trajectories starting on a certain line are confined to it. This is described by the constant of motion Eq. (12). The axes $(h, p)$ are the relative abundances of the focal subtype $(H_1, P_1)$.

In a stochastic model the constant of motion loses this meaning, but it can still be used as an observable to measure the distance of the state to the inner fixed point or the outer border where a type goes extinct. Claussen (2007) uses the average change of this observable to analyse under what conditions the system can be forced to return to the inner fixed point (drift reversal) or whether extinction is inevitable owing to the outward diffusion in a "battle of the sexes" game. In fact, the quantity can be used as an observable also for models that are not neutrally stable, but where $H = 0$ still means extinction and $h, p$ are the two dimensions. Hence, we can use the following approach not only for the Evo process (which is equivalent to the replicator dynamics in a limit) but also the Evo$^+$ process (which is equivalent to the attractive adjusted replicator dynamics). The discrete time versions of these processes allow us to focus on the probabilities only, without having to keep track of time, since $\Delta t = 1$. We now use the average change of this observable to analyse whether the system can be forced to return to the inner fixed point or whether extinction is inevitable due to the outward diffusion.

When a transition occurs, the value of the observable $H$ also changes. For each transition $\mathcal{T}_l(h, p)$ with $l = 1, 2, ..., 9$ and corresponding $\Delta h$ and $\Delta p$ there is a $\Delta H_l(h, p) := H(h + \Delta h, p + \Delta p) - H(h, p)$. By weighing all possible changes in $H$ with the corresponding transition probabilities one receives the average change of the observable for each configuration of host and parasite populations.

$$\overline{\Delta H}(h, p) = \sum_{l=1}^{9} \Delta H_l(h, p)\, \mathcal{T}_l(h, p) \tag{13}$$

This is the average change in $H$ for each coordinate $(h, p)$ in each time step, but the actual extinction time remains to be calculated.

The average diffusion over the whole space would be the integration over $h, p \in [0, 1]$, which is not possible in all generality. It becomes possible when taking the weak selection approximation via a Taylor approximation

$$\overline{\Delta H}_T(h, p) = \overline{\Delta H}(h, p) + \mathcal{O}(w_i^3) \tag{14}$$

around $w_H = w_P = 0$ is up to second order in $w_i$, $i \in \{H, P\}$. After this approximation it is possible to integrate the formula to get a (phase space) average of the average diffusion.

$$\langle\overline{\Delta H}\rangle = \int_0^1 \int_0^1 \overline{\Delta H_T}(h,p)\, dh\, dp \tag{15}$$

For the matching allele infection matrices $M^H = \left(\begin{smallmatrix}0 & 1\\ 1 & 0\end{smallmatrix}\right)$ and $M^P = \left(\begin{smallmatrix}1 & 0\\ 0 & 1\end{smallmatrix}\right)$ in a discrete time Evo or Evo$^+$ process the average diffusion is

$$\langle\overline{\Delta H}\rangle_{dtEvo} = \frac{-100(N_H^2 + N_P^2 - 1) - 4N_H N_P w_H w_P}{225 N_H^2 N_P^2} \tag{16}$$

$$\langle\overline{\Delta H}\rangle_{dtEvo^+} = \frac{A + B\, w_H^2 - C\, w_H w_P + D\, w_P^2}{225 N_H^2 N_P^2} \tag{17}$$

with

$$\begin{aligned}
A &= -200(N_H^2 + N_P^2 - 2)\\
B &= 4 - 2N_P^2 + N_H(2N_P^2 - 4)\\
C &= 4 - 4N_P + N_H(4N_P - 4)\\
D &= 4 - 4N_P + N_H^2(2N_P - 2)
\end{aligned}$$

per time step. Now we can find out the time steps it takes to reach a certain value of $H_{finish}$ starting in a certain value of $H_{start}$.

$$t = \frac{H_{finish} - H_{start}}{\langle\overline{\Delta H}\rangle} \tag{18}$$

We start in $H_{start} = 1$, which is the coexistence fixed point with a maximum variability of host and parasite subtypes. The value of $H$ when one subtype dies out is $H_{finish} = 0$.

$$t = \frac{-1}{\langle\overline{\Delta H}\rangle}. \tag{19}$$

This method only works if the average diffusion is negative. For example, with $w_H = w_P = 1$ and $N_H = 250$, $\langle\overline{\Delta H}\rangle_{dtEvo}$ is always negative but $\langle\overline{\Delta H}\rangle_{dtEvo^+}$ becomes positive for $N_P \geq 85$. The stronger the selection intensity in the dtEvo$^+$ process, the more the pull towards the coexistence (attractive) fixed point. For the dtEvo, the selection intensity has hardly any influence (there are only terms in $\frac{1}{N_i^2}$, whereas in the dtEvo$^+$ there are terms in $\frac{1}{N_i}$).
One has to be careful to use this formula since it was derived in a weak selection limit and large $N$. Yet the simulations present results for small $N$ and relatively large $w$. However, we see that the order of magnitude is the same as in the simulated extinction times above. Yet the logarithmic scale, although convenient to represent the different orders of magnitude, may be misleading.

### S6 Exact sojourn times

The constant population size processes with discrete time (dtEvo$^+$ and dtEvo) allow for a description of transitions as probabilities in each discrete time step $\Delta t = 1$. For each state with index $k \in \{1, 2, ...(N_H + 1)(N_P + 1)\}$ in the two-dimensional state space $H_1 \in \{0, 1, ..., N_H\} \times P_1 \in \{0, 1, ..., N_P\}$ there is a probability of transitioning to every other state (most of these probabilities are zero, because only one step is taken in state space for each time step, see also Table S1). This transition matrix $\Pi \in [0,1]^{(N_H+1)(N_P+1)}$ can be ordered into a block matrix containing one block for transient states $Q$, one block for transitions from transient states to absorbing states $R$, and one identity matrix block for absorbing states $I$. Then the transition matrix is

$$\Pi = \begin{pmatrix} Q & R \\ 0 & I \end{pmatrix}, \tag{20}$$

and the recursion equation

$$\rho(t+1) = \rho(t)\,\Pi, \tag{21}$$

for the probability density $\rho(t) = (\rho_k)_k$ over all states $k \in \{1, 2, .., (N_H+1)(N_P+1)\}$. Grinstead and Snell (2012) proved that the time spent in the inner states, which are part of the matrix $Q$, depending on the initial state is an entry in $(Q-I)^{-1}$. Hindersin et al. (2016) apply this by solving the equivalent system of equations $(Q-I)x = -1$ for $x$. We have applied this numerical approach to the inner states $H_1 \in \{1, 2, ..., N_H - 1\} \times P_1 \in \{1, 2, ..., N_P - 1\}$ which then gives the exact average time to extinction of one of the subtypes in one of the populations.

### S7 Literature overview

In the main text, Table 1 summarises some mathematical models on host-parasite dynamics found in the literature. We have based the choice of our models on the visibility of the studies, the relatedness and comparability to our work and the availability of the information found in the publications, that is necessary to complete the table. We have aimed at focussing on explicit host-parasite models (HP), that build on the classic Lotka-Volterra or Rosenzweig-MacArthur models or the evolutionary game theory literature. There are many such publications from before the year 2000 (Schaffer and Rosenzweig, 1978; Seger, 1988; Nee, 1989, and more), but we aimed at comparing predominantly the more recent literature. However, many models are host-focussed, such as susceptible-infected (SI) epidemiological models. Many host centred models include parasite genotype frequencies and infection probabilities, but the host dynamics are more complex than the parasite dynamics, and the focus is often on host evolution or the maintenance of sexual reproduction in the host. We have briefly discussed these studies, yet we are predominantly interested in co-evolution models, where parasites play a similarly important role (HP models). Furthermore, we have also included some studies that are concerned with slightly less related topics to show the diversity in assumptions and authors that this field provides. Some of these models include stochasticity and a changing population size (spatial structure model of Boots and Sasaki (1999)) while others include stochasticity but not a variable population size (model on special host or parasite behaviour Abou Chakra et al. (2014)) and others include a variable population size but are deterministic (non-host-parasite Red Queen dynamics Bonachela et al. (2017)). This is, naturally, a subjective approach, and we have not excluded any publications for belligerent reasons. We also wish to caution, that this study is not a systematic literature review and we do not make the claim to have collected all relevant publication in this extensive field.
